# Myosin light chain proteins cooperatively promote sarcomere growth in fast-twitch muscle

**DOI:** 10.1101/2024.09.18.613721

**Authors:** Tayo E Adekeye, Troy E Hupper, Teresa E Easterbrooks, Emily M Teets, Emily A Tomak, Sadie L Waterman, Kailee A Sprague, Angelina White, Maddison L Coffin, Daniel K Tanaka, Sabrina M Varga, Mason T Soares, Sarah J Shepherd, Jared D Austin, Dmitrii Krivorotko, Emma S Perry, Sharon L Amacher, Joshua B Kelley, Jared C Talbot

## Abstract

Muscle becomes stronger by expanding sarcomeres from the cell’s edge to its center. Here we investigate how Mylpf (Myosin Light Chain Phosphorylatable Fast) abundance impacts sarcomere growth. The two zebrafish Mylpf genes (*mylpfa* and *mylpfb*) are exclusively expressed in fast-twitch muscle, with *mylpfa* expressed more abundantly than *mylpfb*. Mutations in both genes cause full mRNA and protein loss, and loss of sarcomere formation. The degree of sarcomere reduction in fast-twitch muscle is predicted by Mylpf protein dosage using a high-slope Hill equation. The sarcomere defect may be driven by thick filament localization, which is impaired in *mylpfa^-/-^*and obliterated in *mylpfa^-/-^;mylpfb^-/-^*. Slow-twitch muscle is spared, becoming larger and more active in the mutants. The *mylpfa^-/-^* defects can be rescued by Mylpfa-GFP, Mylpfb-GFP, and by human MYLPF-GFP with equivalent efficiency, but not by a MYLPF protein variant that causes Distal Arthrogryposis. These effects indicate that Mylpf dosage strengthens muscle by mediating myofilament localization, and thereby sarcomere and myofibril growth.

## Introduction

Sarcomeres grow from the edge to the center of muscle cells (the ‘width axis’) and this growth is what strengthens the muscle. Despite the importance of the sarcomere’s width to muscle strength, it remains unclear how its growth is regulated. One way to control a sarcomere’s growth is to regulate the abundance of its structural proteins. Overall protein levels (’protein dose’) are influenced by many factors including a gene’s copy number (’gene dosage’), the amount of steady-state transcript produced by each gene (’mRNA abundance’), the measured amount of protein encoded by these genes (’protein abundance’), the activity of each encoded protein, and the protein’s localization within a cell. In biochemical settings, the relationship between protein dose and its effect can be modeled using four-parameter logistic regressions, such as the Hill equation^1^. The same equation is often used to assess drug efficiency *in vivo*, though it is not commonly applied to investigate how native protein impacts muscle formation. In this study, we investigate how the abundance of one key sarcomeric component, Mylpf (Myosin Light Chain Phosphorylatable Fast), impacts sarcomere formation. We find that the protein is essential to an early step in sarcomere formation and its effects can be efficiently modeled as a Hill response.

Sarcomere growth is best understood in the context of its organelle, the myofibril, which contains chains of sarcomeres that stretch along the length axis of muscle. Myofibril formation is initially directed by actin-rich thin filaments aligned along the edge of the muscle cell ^2–4^. These thin filaments contain F-actin strands bundled with nebulin and linked to actinin-rich Z-disks, producing I-Z-I bodies. The ends of I-Z-I bodies are temporarily connected to one another by non-muscle myosin to form pre-myofibrils, which act as a template for further growth ^4–6^. Growth is also facilitated by integration of enormous contractile proteins titin and nebulin, which set the sarcomere’s length ^7^. Once pre-myofibrils have formed, the non-muscle myosin is quickly replaced by contractile myosin heavy chain (MyHC), bundled into double-headed thick filaments. These added thick filaments increase tension across the muscle cell, leading to myofibrillar growth ^8–10^. Myofibrils grow to only around 2 µm wide, but multilayered arrays of myofibrils (myofibril bundles) continue this growth until they extend deep into the central cytoplasm ^6,10,11^. The myofibril bundles in vertebrate muscle are aligned and tightly interlinked along the width axis of a sarcomere, such that the sarcomere and myofibril bundle have matched width (Figure 1A). In this work, we refer to the sarcomere when examining localization of thick/thin filament components and the myofibril bundle when viewing the structure at a larger scale.

**Figure 1:**
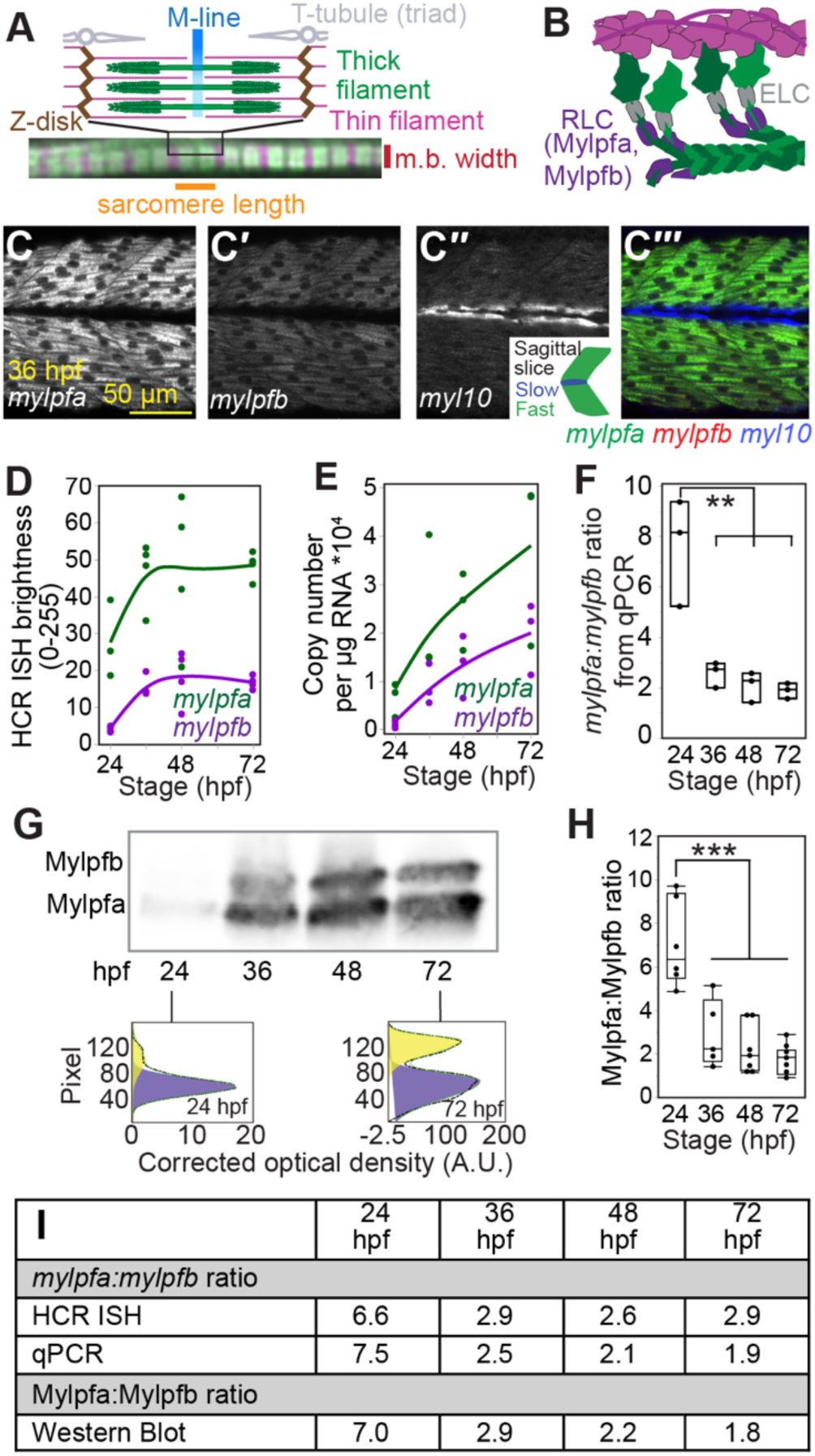
The *mylpfa* gene is expressed more abundantly than *mylpfb* in fast-twitch muscle. **(A)** Images illustrating myofibril and sarcomere structure. The m.b. width represents axis of measurements for myofibril bundle width. **(B)** Illustration of Mylpf localization within a thick filament. **(C-C’’’)** *In situ* hybridization chain reaction (HCR ISH) imaged in somites over the mid-yolk tube of a 36 hpf embryo. Shown as a single channel for *mylpfa* (C), *mylpfb* (C’) or the slow muscle marker *myl10* (C’’), and as a merged image (C’’’). **(D)** Plot showing the brightness of *mylpfa* and *mylpfb* in the HCR ISH images. **(E)** Copy number of *mylpfa* and *mylpfb* per microgram of total RNA at 24, 36, 48, and 72 hpf as measured by qPCR and normalized using a standard curve–based normalization method. **(F)** The ratio of *mylpfa* to *mylpfb* transcript abundance calculated from the absolute copy numbers shown in (E). **(G)** Image of a western blot showing Mylpfa and Mylpfb protein abundance at these timepoints. Examples of Mylpfa and Mylpfb bands quantified in GelBox are shown beneath the image. **(H)** Ratio of Mylpfa to Mylpfb band intensity in western blot. **(I)** Tabular summary of means from D, F, H. The points represent statistical N, either pools of animals (E, F, H), or individual animals (D). Significance threshold determined by Tukey-Kramer comparison after one-way ANOVA; ** P<0.01, ***P<0.001.

The class II MyHC in thick filaments is stabilized close to its force-generating head by a Regulatory Light Chain (RLC) and an Essential Light Chain (ELC) ^12,13^ (Figure 1B). These light chains regulate myosin movement and force generation but do not consume ATP nor produce force on their own ^12^. Point mutations in the RLCs can reduce myosin step size ^14^, and outright removal of these light chains *in vitro* causes MyHC aggregation and partially reduces MyHC activity ^15,16^. Light chains also have critical roles *in vivo,* as first shown in the *Drosophila* RLC mutant, which lacks skeletal muscle^17^. Similarly, the ELC gene *Myl1* is required for normal myogenesis in chicken, zebrafish, mouse, and human ^18–20^. Because the primary effect of these light chains is to regulate and stabilize MyHC, we hypothesize that myosin light chain proteins influence the rate of cytoskeletal organization, leading to myofibril growth.

The only RLC with prominent expression in embryonic muscle fibers and postnatal fast-twitch skeletal muscle of mouse is Mylpf ^21^. All skeletal muscle fibers are absent from the mouse *Mylpf* knockout at birth ^22^. This defect is so dramatic that it is unclear what phase of myogenesis is disrupted by the mutation. *MYLPF* function is also critical for human development. Missense alleles in *MYLPF* cause Distal Arthrogryposis (DA), a congenital musculoskeletal disease characterized by inherited distal limb contractures^23^. DA is often caused by mutations in genes that encode contractile proteins ^24,25^. However, neither muscle strength nor structure were examined in human patients with the *MYLPF* variants, and the impact of those variants has not been tested in animal models. Therefore, the impact of *MYLPF* missense alleles on muscle formation remains unresolved.

Muscle formation in vertebrates begins in segments of mesoderm called somites, which eventually produce most of the muscles in the body. Zebrafish somites produce two fiber types, fast-twitch and slow-twitch, within the first day post fertilization (dpf) ^26^. Whereas slow-twitch fibers contract and fatigue slowly, the zebrafish fast-twitch fibers contract with such speed and power that they strain muscle cells nearly to the point of snapping ^27,28^. By 24 hours post fertilization (hpf), zebrafish fast-twitch muscle fibers are positioned medial to a thin layer of slow-twitch fibers ^26^, enabling us to identify fiber types by position in addition to molecular markers of fiber type. Embryonic muscle fibers grow rapidly, more than doubling in size by the time of hatching (3 dpf) ^29^. Hatched larvae continue growing while subsisting on their yolk until 6 dpf, enabling study of early larval growth independent of feeding. In zebrafish, Mylpf gene function is distributed across two paralogs, *mylpfa* and *mylpfb* ^30–32^. We previously showed that the zebrafish *mylpfa^-/-^*mutant can form multinucleate muscle cells but have weakened muscle that eventually deteriorates ^33^. This mutant has severe physiological defects including impaired escape response and 90% loss in muscle strength, suggesting that there may be effects on the myofibril itself ^33^. However, the impact of *mylpfa* on sarcomere formation was untested and the requirements for *mylpfb* function in embryonic development remained unexplored.

Here, we investigate the effect of Mylpf activity on sarcomere growth along the width axis of muscle fibers. We generated a frameshifting *mylpfb^-/-^* mutant, which has no overt muscle defect alone, but dramatically enhances the *mylpfa^-/-^* mutant phenotype. The *mylpfa^-/-^* mutant has a severe reduction in myofilament overlap in fast-twitch muscle, leading to impaired sarcomere growth and thereby less myofibril bundle growth within that fiber type. Actin-myosin colocalization fails entirely in the fast-twitch muscle of the *mylpfa^-/-^;mylpfb^-/-^*double mutant, causing a complete absence of fast-twitch contractile structures. Both mutants are null for protein production due to nonsense mediated decay. We find no evidence for compensation for Mylpf loss of function among *mylpfa* and *mylpfb* genes, though the mutant animals do compensate in slow-twitch myofibril bundle size and slow movement. Transgenic expression experiments in the *mylpfa^-/-^*mutant suggest that function and efficiency is conserved among the Mylpfa, Mylpfb, and human MYLPF proteins. By contrast, a transgene that expresses a protein variant found in Distal Arthrogryposis patients does not rescue muscle structure. Dose analysis in knockout and transgenic animals suggests that Mylpf binding has a cooperative effect on myofibril bundle growth, with initial binding enabling a more-than-additive subsequent increase until saturation is approached. Together, these findings show a critical impact of Mylpf dosage on sarcomere and myofibril production during early fast-twitch myogenesis.

## Results

### Mylpfa is more abundant than Mylpfb during embryonic development

Although *mylpfa* and *mylpfb* expression patterns have been described individually ^34^, their overlap and relative abundance have not been clarified. Each gene produces one detectable transcript exclusively in fast-twitch muscle fibers (Figures S1, S2). Expression begins by 20 hpf in medial fast-twitch fibers (Figure S2). Neither gene is expressed in slow-twitch muscle, which expresses a different regulatory light chain, *myl10* (Figure 1C-C”’) ^35^. In both HCR and qPCR, *mylpfa* is expressed more abundantly than *mylpfb*; the ratio between these genes is roughly 7:1 at 1 dpf and diminishes to 2:1 by 48 hpf, largely because of rising *mylpfb* mRNA abundance (Figure 1D-F). These ratios can also be found in publicly available RNA-seq data ^33^. These zebrafish transcripts encode Mylpf proteins that are 94% identical to one another and each is around 80% identical to human MYLPF. Using newly generated antibodies we can distinguish the two proteins on western blot, which differ in size by 2 kilodaltons (kD). The relative protein abundance correlates with transcript levels (Figure 1G-I). In summary, the two zebrafish Mylpf genes are expressed in fast-twitch muscle, producing Mylpfa protein more abundantly than Mylpfb during embryonic stages.

### The *mylpfa^oz30^* and *mylpfb^oz39^* alleles are effectively null

We had previously made an allele of *mylpfa* (*oz30*) ^33^, and here provide evidence that it causes severe sarcomere and myofibril bundle defects (Figure 2A-G). These myofibril bundle defects are also apparent in confocal stack of phalloidin-labeled *mylpfa^-/-^* mutant (Video 1) and in multiple quantification methods (Figures 2I-M S3, S4). Although the *mylpfa^-/-^*mutant has severe myofibril reduction at all timepoints, it forms differentiated somite muscle, which has wild-type size at 2 dpf and a modest growth defect by 6 dpf (Figure 2H). These effects are confirmed by a second allele in the same gene (Figure S5) ^33^. Both *mylpfa* mutants can at least partially form myofibrils in fast-twitch muscle (Figure S5) so we hypothesized that *mylpfb* may also contribute to function. We mirrored the *mylpfa^oz30^* mutant by generating a 5 bp deletion in *mylpfb* (allele *oz39*) that frameshifts the *mylpfb* transcript at AA 78 out of 170 (Figure 3A-C). Using RT-qPCR we find that mRNA levels for *mylpfa* and *mylpfb* are approximately halved in heterozygotes and essentially undetectable in each mutant (Figure 3D-G), suggesting that both mutations lose transcript through nonsense mediated decay. Any protein made by this frameshifting allele should be nonfunctional because the encoded peptide has at most only a short stretch capable of binding to MyHC (Figure 3H). However, we find no Mylpfa protein produced in the *mylpfa^-/-^* mutant nor any Mylpfb in the *mylpfb^-/-^* mutant, and neither protein in the *mylpfa^-/-^;mylpfb^-/-^*double mutant at 48 hpf (Figure 3I-J).

**Figure 2:**
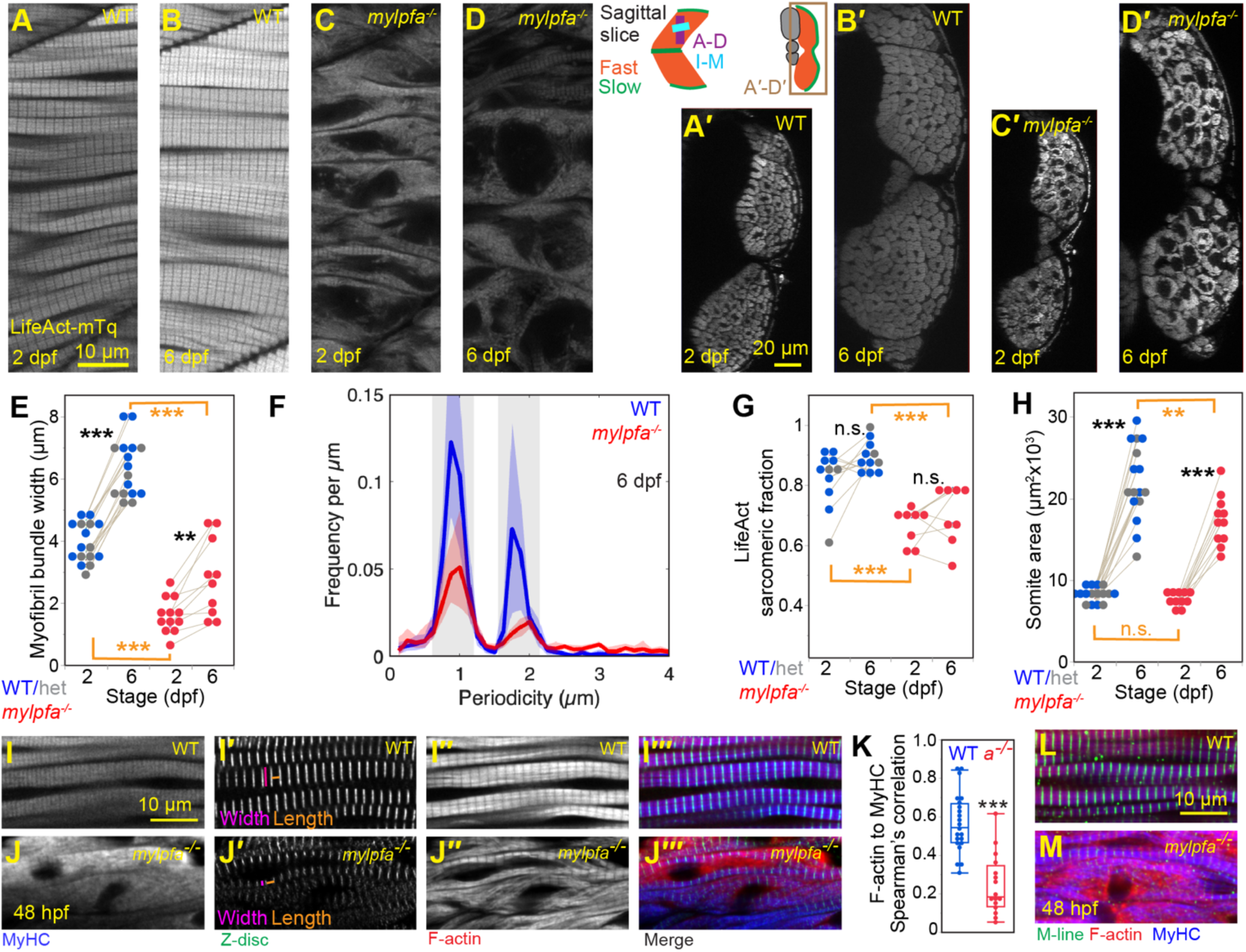
Zebrafish *mylpfa* is necessary for fast-twitch myofibril formation. **(A-D)** Zoomed region of somite muscle, with myofibril bundles marked using *six1b:LifeAct-mTurquoise2* transgene (LifeAct-mTq protein) at 2 dpf and the same animal at 6 dpf in wild-type (A, B) or *mylpfa^-/-^* mutant siblings (C, D). **(A’-D’)** Cross-sectional views of whole somites from the same animals. **(E)** Myofibril bundle widths were measured using the LifeAct images in the sagittal plane. Olive-colored lines connect the same individuals through time. **(F)** Plot of signal periodicity shows that the *mylpfa^-/-^* mutant has less LifeAct periodicity in sarcomeric intervals (gray bars). Lightly colored regions indicate bootstrap confidence intervals. **(G)** This readout is simplified as a ‘sarcomeric fraction’, which shows the fraction of signal in sarcomeric (1.85 µm) or half-sarcomeric (0.9 µm) lengths out of the total periodicity; further explanation is provided in Figure S4. The sarcomeric fraction remains constant through development but differs by genotype. **(H)** Somite area, measured from the cross-sectional views. The *mylpfa^-/-^* mutant somite is smaller than in wild-type or heterozygous siblings at 6 dpf, though not at 2 dpf. **(I-J’’’)** Fast muscle myofibers, labeled at 48 hpf, showing co-label for MyHC (A4.1025), the Z-disk protein Actinin (A7732), and F-actin (phalloidin), shown as single channel or overlays. The axes of width and length measurements are shown in I’, J’. A representative confocal stack of this type of label is shown in Video 1. **(K)** The correlation between F-actin and MyHC label is reduced in the *mylpfa^-/-^* mutant, as assessed using Spearman’s rank correlation coefficient (Spearman’s correlation). **(L, M)** A similar pattern is seen in animals labeled for the M-line protein Myomesin (mMac), F-actin, and MyHC. Scalebar in A is for A-D, in A’ is for A’-D’, in I is for I-J’’’, in L is for L, M. Each point shown represents the average of measurements taken within one image of one animal. Significance thresholds for multiple comparisons determined by Tukey-Kramer HSD comparisons after one-way ANOVA (E, F, H); pairwise comparisons (K) use the Kruskal-Wallis exact test. Not significant (n.s.) is P>0.05, * P<0.05, ** P<0.01, *** P<0.001.

**Figure 3:**
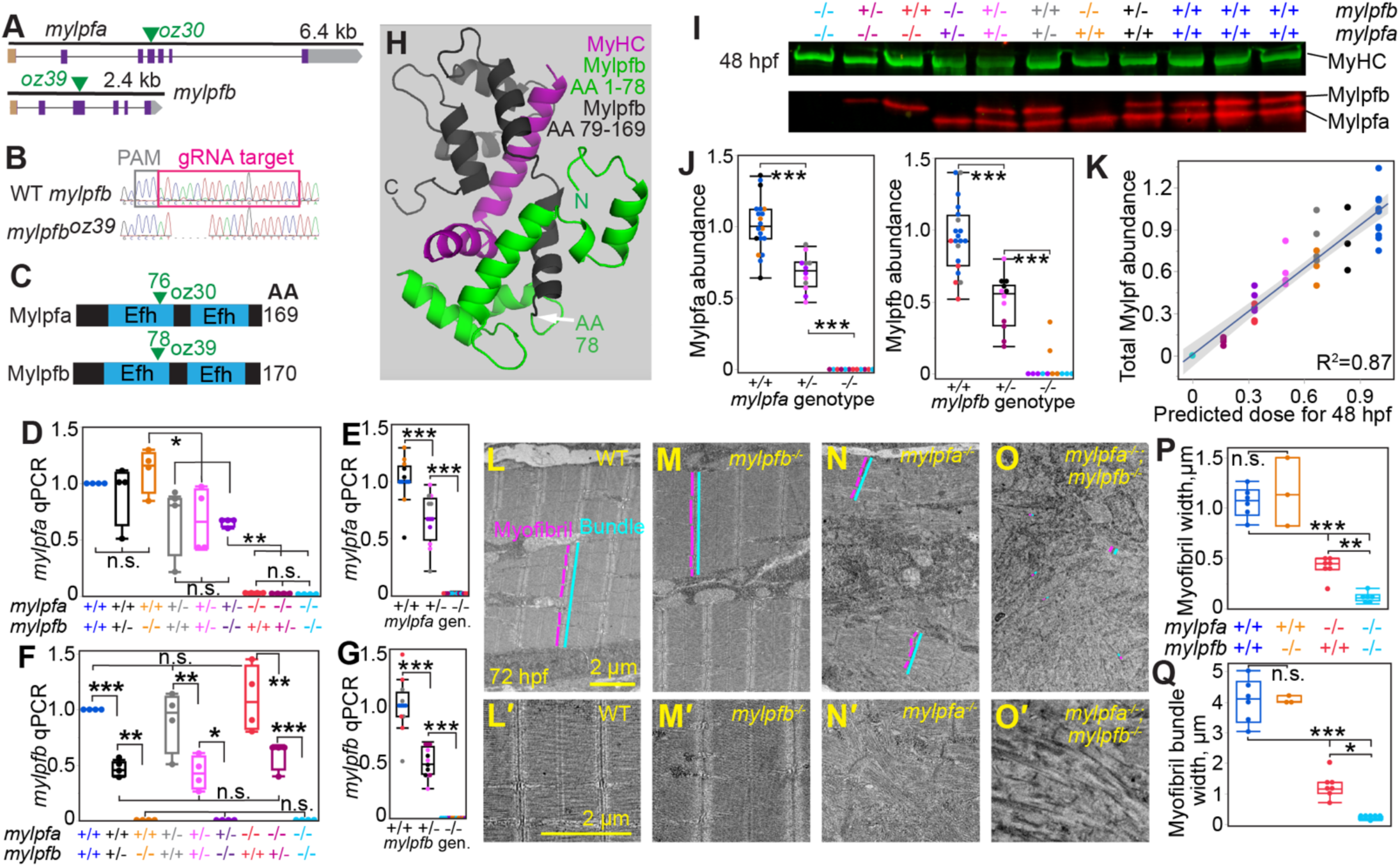
Mutations in *mylpfa* and *mylpfb* cause strong loss of function. **(A)** Illustration of *mylpfa* and *mylpfb* gene structure. Color code: 5’ UTR (brown), coding sequence (purple), 3’ UTR (gray), and frameshift locations (green arrowheads). **(B)** Chromatogram showing the gRNA target in wild-type sequence (top) and the 5 bp *mylpfb^oz39^*lesion sequenced from a homozygous mutant (bottom). **(C)** Illustration of Mylpfa and Mylpfb proteins, with arrowheads marking frameshift locations. **(D-E)** Quantitative RT-PCR results from the *mylpfa;mylpfb* mutant series showing *mylpfa* expression per genotype (gen.) normalized to the average *mylpfa* abundance in *mylpfa^+/+^* genotypes. Result shown as individual genotypes (D) or as a merger of *mylpfa* genotypes (E). **(F-G)** Quantitative RT-PCR results for *mylpfb* expression, normalized to the average *mylpfb* abundance in *mylpfb^+/+^* genotypes, shown for individual genotypes (F) or as a merger of *mylpfb* genotypes (G). **(H)** Predicted protein structure generated using Robetta. Color code indicates the Mylpf peptides lost after *mylpfb^oz39^* frameshift (black), those retained even in the mutant (green), and the MyHC (purple). **(I)** Western blot showing MyHC, Mylpfb, and Mylpfa bands from individual animals across the nine genotypes of *mylpfa^+/-^;mylpfb^+/-^* incross. Three wild-type controls are loaded to increase sampling of the wild-type mean. **(J)** Plot showing the abundance of each protein in animals wild-type, heterozygous, or mutant for that gene. **(K)** Plot showing a strong correspondence between combined Mylpfa and Mylpfb band intensity and predicted dosage using a 2 to 1 ratio between Mylpfa and Mylpfb (Table S1). **(L-O)** TEM images of fast-twitch muscle showing myofibrils in wild-type and their defects in the single and double mutants. Lines illustrate measurements of muscle structure. **(L’-O’)** Higher magnification views of different animals. **(P)** Measurements made between triads, showing myofibril width. **(Q)** Measurements of myofibril bundle widths. Each point (statistical N) in J, K, P, Q represent individual animals; in D-G, each dot represents a pool of three animals. Color code below D is for D-G, I-K, P-Q. Significance thresholds are determined by Tukey-Kramer comparison after one-way ANOVA; n.s. P>0.1, * P<0.05, ** P<0.01, *** P<0.001. Scalebar in L is for L-O, in L’ is for L’-O’.

Examination of all nine genotypes in western blot shows that overall Mylpf dosage (Mylpfa & Mylpfb) corresponds closely to predictions made using the 2:1 dosage seen at 48 hpf, suggesting that there is little intragenic compensation (Figure 3K, Table S1). For instance, the *mylpfa^+/-^* and *mylpfb^+/-^* heterozygotes express around half the normal protein dose (67% for *mylpfa^+/-^*, 49% for *mylpfb^+/-^,* Figure 3J). These findings indicate that the mutants used in this study are essentially null for the single product of the *mylpfa* and *mylpfb* genes, respectively.

### Together, *mylpfa* and *mylpfb* are essential to sarcomere formation

Even though the Mylpfb protein sequence is very similar to Mylpfa, the two mutants have strikingly different phenotypes. The *mylpfb^-/-^*mutant looks entirely wild-type on the electron microscope (Figure 3L-M’) and in every other assay we have tested thus far. By contrast, the *mylpfa^-/-^* mutant has narrowed myofibrils, with poor myofilament alignment on the sarcomere level (Figure 3N, N’). In contrast to the partial defects seen in the *mylpfa^-/-^* mutant, the *mylpfa^-/-^;mylpfb^-/-^* double mutant lacks sarcomeres entirely (Figure 3O, O’). This double mutant shows a catastrophic loss of striation in fast-twitch muscle, more severe than either of the single mutants, suggesting that both genes contribute to muscle structure (Figure 3O-Q). In this double mutant, the fast-twitch muscle shows no true sarcomeres or myofibrils, only a scattering of thick filaments and I-Z-I bodies (Figure 3O, O’). We observe equivalent effects on myofibril width (measured between membrane triads) and myofibril bundle size (measured across interconnected myofibrils) (Figure 3P, Q). Even in the double mutant, somite muscle is present and the abundance of contractile MyHC is unchanged (Figure 3I, P>0.3 in ANOVA), suggesting that muscle differentiation proceeds despite the loss of organized myofibrils. These findings suggest that the two Mylpf genes are collectively essential to sarcomere formation and therefore to myofibril bundle formation, with *mylpfa* playing a larger role.

### The *mylpf* genes are essential to myofilament colocalization

Because Mylpf binds to MyHC, we hypothesized that the sarcomere defect may arise through an initial failure in MyHC localization, independent of initial F-actin ordering. We examined muscle structure in Mylpf mutants at an early stage of myofibril bundling (26 hpf). We find profound organizational defects in fast-twitch muscles, but not slow-twitch fibers (Figure 4A-D). The *mylpfa^-/-^* mutant fails to properly localize MyHC to the fast-twitch muscle cell’s periphery where pre-myofibrils are forming, with stronger defects in the *mylpfa^-/-^;mylpfb^-/-^*double mutant (Figure 4C-D’). Consistent with this localization defect, the correlation between F-actin and MyHC label is reduced in the *mylpfa^-/-^*mutant and worsened in the *mylpfa^-/-^;mylpfb^-/-^* double mutant, to a degree that parallels reductions in myofibril bundle width (Figure 4E-F). The confocal images suggested that sarcomere and myofibril formation may fail in the mutant because F-actin and MyHC are found in different portions of the cell. At this early stage, the pre-myofibrils form rings around cells even in the *mylpfa^-/-^;mylpfb^-/-^* double mutant (Figure 4A-D), consistent with findings that actin localization precedes thick filament localization ^36^. Using these rings of F-actin label, we marked cell boundaries and then measured how far each marker spreads from that boundary to the cell center in fast-twitch muscle cells (Figure 4G). By contrast, each genotype has a similar distribution of F-actin and myonuclei along this cell edge-center axis (Figure 4H, I). MyHC is also localized at cell peripheries in the wild-type embryo, but it spreads evenly through the cytoplasm of the *mylpfa^-/-^;mylpfb^-/-^* double mutant (Figure 4J). These findings suggest that Mylpfa and Mylpfb together are essential to localize MyHC to the fast muscle cell’s periphery where the thick filaments interdigitate with pre-myofibrils to initiate myofibril growth (Figure 4K).

**Figure 4:**
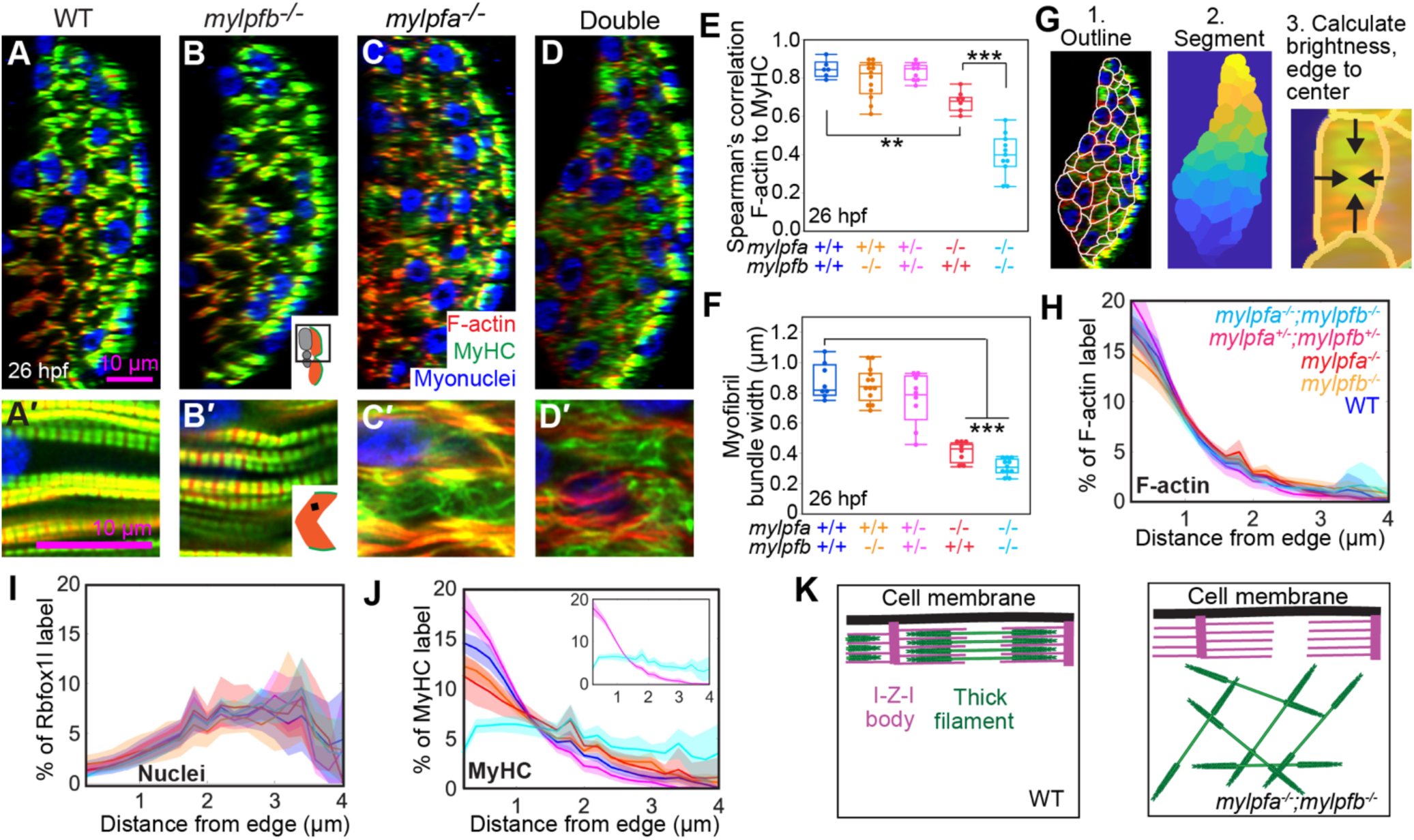
*mylpfa* and *mylpfb* are required for myosin heavy chain localization in fast-twitch muscle of young myofibers. **(A-D)** Immunolabel of fast muscle fibers at 26 hpf shown as orthogonal views of Z-stacks of wild-type (A), *mylpfb^-/-^* (B), *mylpfa^-/-^* (C), and *mylpfa^-/-^;mylpfb^-/-^*double mutant zebrafish (D). **(A’-D’)** Zoomed sagittal views of the same animals. **(E)** Box plots showing Spearman’s correlation between the F-actin and MyHC label for the genotypes examined. **(F)** Plots showing the width of myofibril bundles in each genotype. **(G)** Overview of protocol to measure image brightness from edge-to-center. The fast-twitch muscle cells are (1) manually outlined, (2) segmented using these outlines, then (3) the outlines are used to determine edges when calculating the brightness of each channel from edge to center of the drawn cells. **(H-J)** Localization at 26 hpf of the F-actin marker phalloidin (H), the muscle nuclei label anti-Rbfox1l (I), and MyHC antibody A4.1025 (J). Data is plotted as percent of total image brightness from edge to center of segments, with bootstrapped 95% confidence intervals in shaded lines. Inset in J shows the stark difference between MyHC localization in the *mylpfa^+/-^;mylpfb^+/-^* double heterozygote and the *mylpfa^-/-^;mylpfb^-/-^* double mutant, unobscured by other genotypes. Color code in H is for H-J. **(K)** Illustration of the proposed localization defect. Points in E-F represent measurements from individual animals. Measurements for E-J were taken on 5 wild-type, 8 *mylpfb^-/-^*, 9 *mylpfa^+/-^;mylpfb^+/-^*, 5 *mylpfa^-/-^*, and 7 *mylpfa^-/-^;mylpfb^-/-^* double mutants. Significance thresholds are determined by Bootstrap comparison (H-J) or by Tukey-Kramer comparison after one-way ANOVA (E, F); ** P<0.01, *** P<0.001. Scalebar in A is for A-D, in A’ is for A’-D’.

### Mylpf dosage shows a cooperative effect on myofibril phenotypes

The Mylpf phenotypes appear to arise in a dose sensitive fashion. To test how well dosage predicts phenotype, we calculated dose predictions in mutants using the wild-type ratio of Mylpfa and Mylpfb, ranging from zero in completely null *mylpfa^-/-^;mylpfb^-/-^* to a dose of one in WT *mylpfa^+/+^;mylpfb^+/+^*(Table S1). We then fit gene dosage and the associated phenotypic outcomes to the Hill equation (Figure 5A), a standard approach for assessing dose-response relationships, which allows us to find the half-maximal dose and measure cooperativity of the response. The 26 hpf myofibril bundle widths show a close correspondence (R^2^=0.84) to dosage at that stage (7:1, 7 Mylpfa per 1 Mylpfb). We then measured muscle phenotype in animals fixed at 2, 3, and 6 dpf and modeled the dose response using the 2:1 Mylpfa to Mylpfb ratio found at later timepoints (Table S1, Figure 5B-D). At all timepoints we found a sigmoidal gradient of fast-twitch myofibril bundle widths from the double mutant (narrowest/none) to wild-type (widest) genotypes, including the 6 dpf timepoint where we analyzed all nine genotypes of the *mylpfa;mylpfb* mutant series (Figure 5A-E). The degree to which myofibrils fill muscle changes more dramatically than the somite muscle’s size (Figure 5D-F). The changes in myofibril bundle width can be explained by the underlying loss of localization between F-actin and MyHC, which by 48 hpf both localize poorly with one another and to the sarcomere in the *mylpfa^-/-^;mylpfb^-/-^* double mutant (Figure 5G-J). Across all genotypes, timepoints, and assays, mRNA and protein dose are better predictors of phenotype than gene dose (Table S2). Early Mylpf dosage has long-lasting impact, since the four genotypes with lowest Mylpf dose in embryos are inviable by adulthood, whereas animals showing high embryonic Mylpf levels (such as the *mylpfb^-/-^* mutant) can grow to a normal adult size (Figure S6). The response curves show a half-maximal protein dose (EC50) of 0.38 on average, suggesting that the Mylpf protein is over halfway to its maximal effect in the double heterozygote (*mylpfa^+/-^;mylpfb^+/-^*, Mylpf dose=0.5). Furthermore, by 48 hpf the response curves consistently have high Hill slopes (>4), indicating that Mylpf levels have cooperative effects in the muscle (Figure 5K). This cooperativity suggests that the presence of Mylpf enhances the impact of additional Mylpf levels on sarcomere growth and thereby on myofibril bundle size, with lasting effects on animal health.

**Figure 5:**
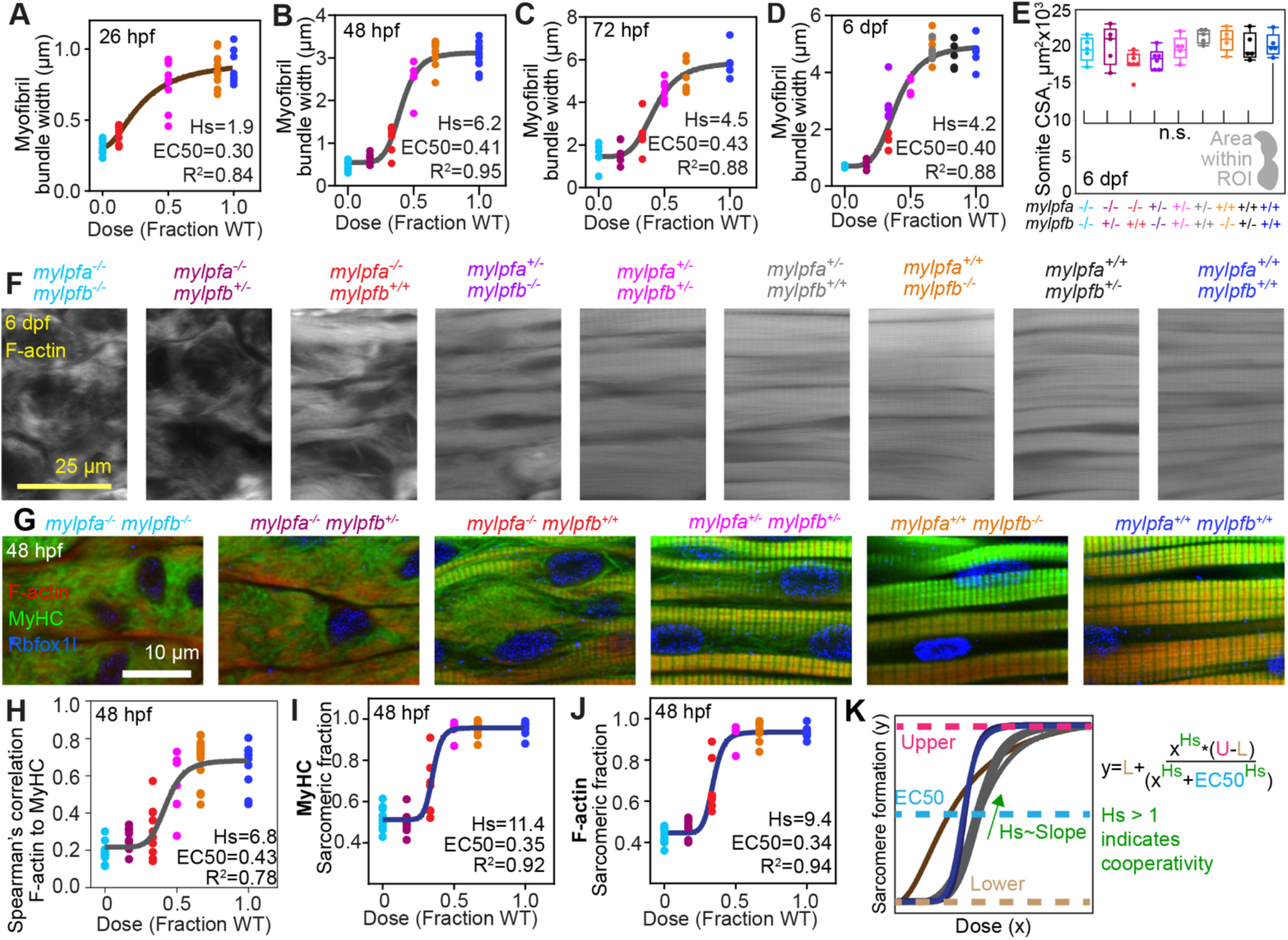
Levels of myofibril formation correspond to dosages predicted by *mylpfa* and *mylpfb* loss of function. **(A)** Scatterplot of myofibril bundle widths in all genotypes tested at 26 hpf, with the X-axis representing Mylpf dosage as predicted by a 7:1 ratio between Mylpfa and Mylpfb. This 26 hpf width data is also shown in Figure 4F, for Tukey-Kramer comparisons. **(B-D)** Scatterplots of myofibril bundle widths in all genotypes tested at 48 hpf, 72 hpf, and 6 dpf, with the X-axis representing Mylpf doses predicted by a 2:1 ratio between Mylpfa and Mylpfb. **(E)** Cross-sectional area measurements of the same somites at 6 dpf. The somite sizes do not differ significantly across genotypes. **(F)** Examples of fast-twitch muscle structure, labeled with phalloidin, in the nine genotypes arising from *mylpfa^+/-^;mylpfb^+/-^* incross, viewed at high magnification. **(G)** Fast-twitch myofibril bundles at 48 hpf, labeled for the myonuclei (Rbfox1l, blue), F-actin (phalloidin, red), and MyHC (A4.1025, green). **(H)** The colocalization between F-actin and MyHC label corresponds well with modeled protein dosage, as measured using Spearman’s correlation, and dose determined with a 2:1 Mylpfa to Mylpfb ratio. **(I-J)** Graph showing how the sarcomeric fraction of MyHC (I) or F-actin (J) corresponds with protein dosage in fast-twitch muscle, again calculated using a 2 to 1 skew. **(K)** Overlay of the 7 regression models with explanation of the Hill equation, scaled to have matching upper and lower limits. Overlay colors in K use regression color code from panels in A (brown), B-D (gray), H (gray), and I, J (blue). Formula abbreviations: Hill slope (Hs), dose of half-maximal effectiveness (EC50), upper asymptote (U), and lower asymptote (L). Each point represents a statistical N, the mean of measurements from one animal. R^2^ and Hs plotted in each panel. The genotypic color code shown above F is used for panels A-E, H-J.

### Slow-twitch muscle compensates for impaired fast-twitch muscle

Zebrafish movement is driven largely by force generated in somite muscle, which is primarily fast-twitch ^26,27^. Consistent with its fast-twitch impairment, our prior work showed that *mylpfa* is essential to muscle strength and fish movement, leading to a severely impaired escape response in the *mylpfa^-/-^* mutant ^33^. In further support, high-speed imaging shows that the sharp flexure driving initial escape response (the ‘C-bend’) is consistently absent from this mutant (Figure 6A-C, Video 2). To learn how behaviors change through time, we assessed voluntary movement by imaging larvae in 25-minute sessions daily from 2 to 6 dpf. Swimming became more frequent in both genotypes through this developmental interval. Surprisingly, by 6 dpf the *mylpfa^-/-^*mutant moves further than their wild-type siblings (Figure 6D). As expected, the *mylpfa^-/-^*mutant cannot swim quickly at any of the timepoints, but by 5 dpf it increases the rate of slow movement compared to its wild-type sibling (Figure 6E-F). To test whether the remaining muscle function came from *mylpfb*, we also examined animals carrying all nine combinations of *mylpfa* and *mylpfb* zygosity at 6 dpf (Figure 6G-J). Consistent with the *mylpfa^-/-^* single mutant, the overall speed is not decreased in the *mylpfa^-/-^*;*mylpfb^-/-^*double mutant, which travels at least as far as wild-type siblings during the imaging period (Figure 6G). However, the highest movement speed is reduced in the *mylpfa^-/-^*animals and also in the *mylpfa^-/-^;mylpfb^-/-^* double mutant, which do not swim faster than 70 mm/sec (Figure 6H), consistent with the escape response defect. The genotypes with severe loss of Mylpf function increase the percent of time that they swim slowly (20-40 mm/sec), enough to compensate for the loss of movement at high speed (>70 mm/sec) (Figures 6I, J, S7). This increased frequency of slow movement may be explained in part by increased slow-twitch myofibril bundle size, which is unchanged across the mutant series at 2 dpf but expands in the severe Mylpf mutants by 6 dpf (Figures 6K-M, S8). We also find increased movement in the *mylpfa^+/-^;mylpfb^+/-^* double heterozygote, which moves faster on average (and therefore further) than wild-type (Figure 6G, I, J Pink), by swimming more often at slow and rapid speeds. This double heterozygote has a modest fast-twitch deficit and no slow-twitch hypertrophy at 6 dpf (Figure 5D, 6M Pink). Together, these findings suggest that slow-twitch muscle utilization and size compensate proportionately with fast-twitch muscle impairment (Figure 6N).

**Figure 6:**
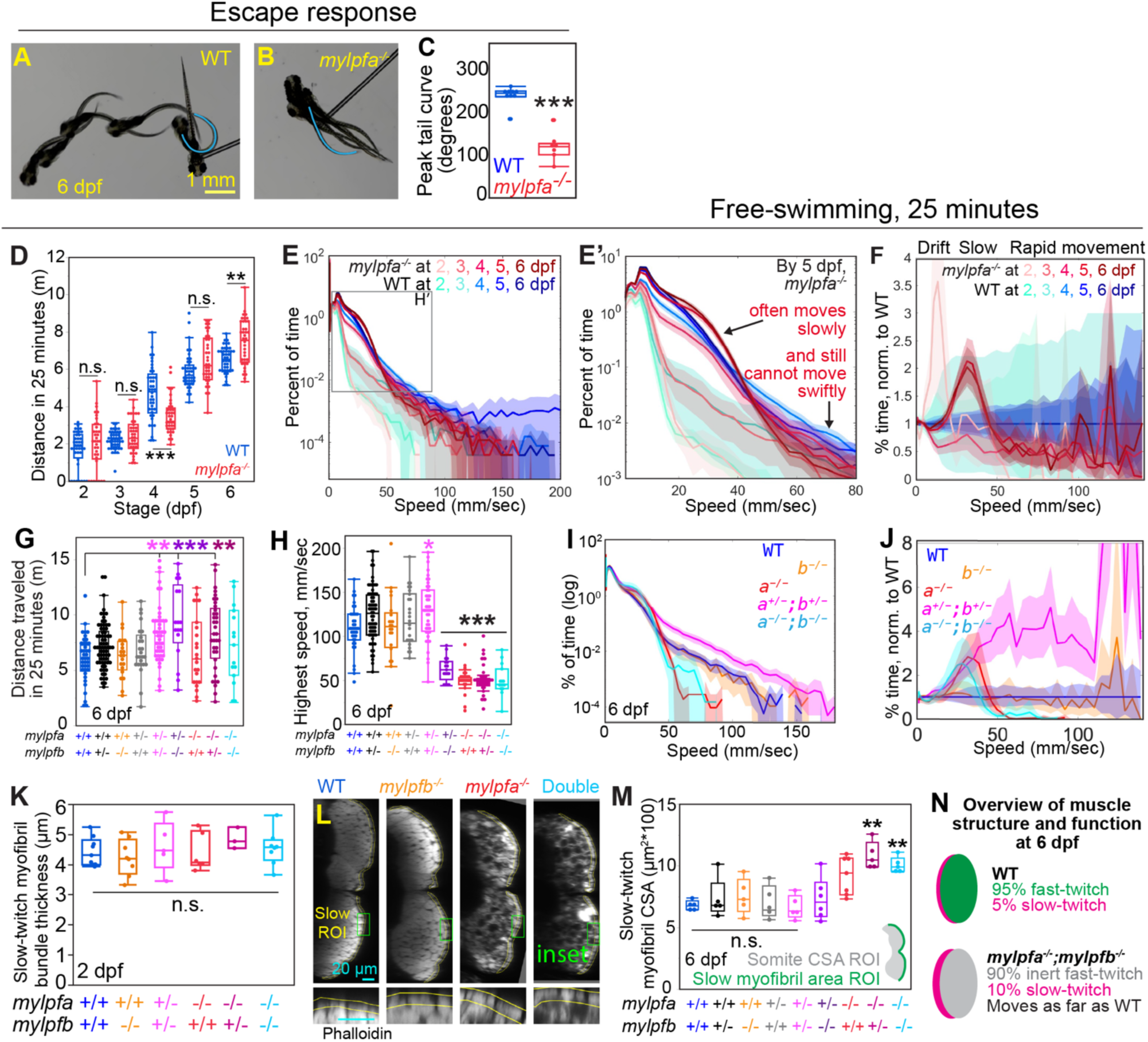
The *mylpfa^-/-^* and the *mylpfa^-/-^; mylpfb^-/-^* double mutant animals do not swim at high speed but increase the frequency of slower movements. **(A-C)** Escape response in wild-type or in *mylpfa^-/-^*animals, showing images captured 1/60th of a second apart (A, B). Blue line shows maximal tail bend, quantified in (C)**. (D)** Box plot of distances traveled when the same fish were imaged 25-minutes per day from 2 to 6 dpf. **(E-F)** A logarithmic plot of the proportion of time that select genotypes swim at different speeds, with bootstrap confidence intervals shown for each genotype (E). The inset region highlights differences in the *mylpfa^-/-^* mutant behaviors that become apparent by 5 dpf (E’). These differences are more striking when normalized to wild-type siblings (F). **(G, H)** Box plots show total distance traveled (G) or highest movement speed (H) of the *mylpfa;mylpfb* mutant series at 6 dpf. **(I)** A logarithmic plot of the proportion of time that select genotypes swim at different speeds, with bootstrap confidence intervals shown for each genotype. **(J)** The same data is shown after normalization to wild-type behaviors, plotted on a linear scale. **(K)** Quantification of slow-twitch myofibril size. **(L)** By 6 dpf the slow-twitch myofibrils bundles are wider in the non-transgenic mutant than their wild-type siblings, seen in cross sectional views, with inset showing the measured region. **(M)** Quantification of the myofibril bundle CSA within the selected slow-twitch regions. No significant difference is seen among high-dose Mylpf genotypes, but at low dose, there is a significant increase in myofibril bundle size, only within slow-twitch fibers. **(N)** Illustration summarizing the effects of Mylpf loss on slow and fast-twitch muscle. Data in (K-M) comes from datasets examined in Figure 5 for other measures. Scalebar in A is for A-B. Significance is determined using the Kruskal-Wallis exact test (C), comparison of bootstrap confidence intervals (E-F, I-J) or Tukey-Kramer comparisons after one-way ANOVA (D, G, H, K, M). Significance shown in comparison to wild-type siblings, with n.s. of P>0.1, * P<0.05, ** P<0.01 and *** P<0.001. Genotypic color code is matched across panels G-J, K, and M.

### Mylpfa-GFP and Mylpfb-GFP restore *mylpfa^-/-^*muscle in proportion to dosage

To further test parallels between Mylpf expression levels and myofibril width, we made a *mylpfa-GFP* transgenic line, *Tg(mylpfa:mylpfa-GFP)mai102,* and found that it can restore myofibril formation in the *mylpfa^-/-^* mutant (Figure 7A-H). The degree of rescue (Figure 7G, H) and the brightness of Mylpfa-GFP (Figure 7B) expression varies among individuals, and we reasoned that Mylpf dosage may impact the degree of rescue. This transgene uses the *mylpfa* promoter and drives GFP-tagged Mylpfa protein at around the same average abundance as the native wild-type Mylpfa (Figure 7C, D). The native Mylpfa protein also varies in expression, though not as widely as Mylpfa-GFP (Figure 7D). The transgene’s fluorescent signal corresponds with protein abundance as measured in western blot, so we used fluorescence as a proxy for protein levels (Figure 7E). Both the myofibril bundle width and the sarcomeric fraction for F-actin correlate closely with Mylpfa-GFP abundance within the *mylpfa^-/-^* genotype; the dimmer fish have widths that are similar to their non-transgenic siblings and the brightest fish overlap with the wild-type range (Figure 7I, J). However, in zebrafish that are wild-type for *mylpfa*, the transgene causes no significant improvement in fast-twitch myofibril bundle size. We found a similar restoration of myofibril bundle formation in the *mylpfa^-/-^*mutant animals that express Mylpfb-GFP from transgene *Tg(mylpfa:mylpfb-GFP)mai103* (Figures 7F-J, S9). The equivalent rescue by Mylpfb-GFP supports our proposal that gene expression levels dictate the difference in *mylpfa* and *mylpfb* requirements, rather than protein sequence. Likewise, we found comparable effects when we examined the effects of Mylpfa-GFP at 6 dpf (Figure S10). Not only does the transgene increase the fast-twitch myofibril bundle width in *mylpfa^-/-^*mutant, but it also suppresses the compensatory hypertrophy of slow-twitch myofibrils in the same animals, restoring both fiber types to their wild-type state (Figures 7, S10). Together, these findings support a model where Mylpfa and Mylpfb are required in proportion to their wild-type dosage, but neither is sufficient to promote excess myofibril bundle growth in a wild-type animal.

**Figure 7:**
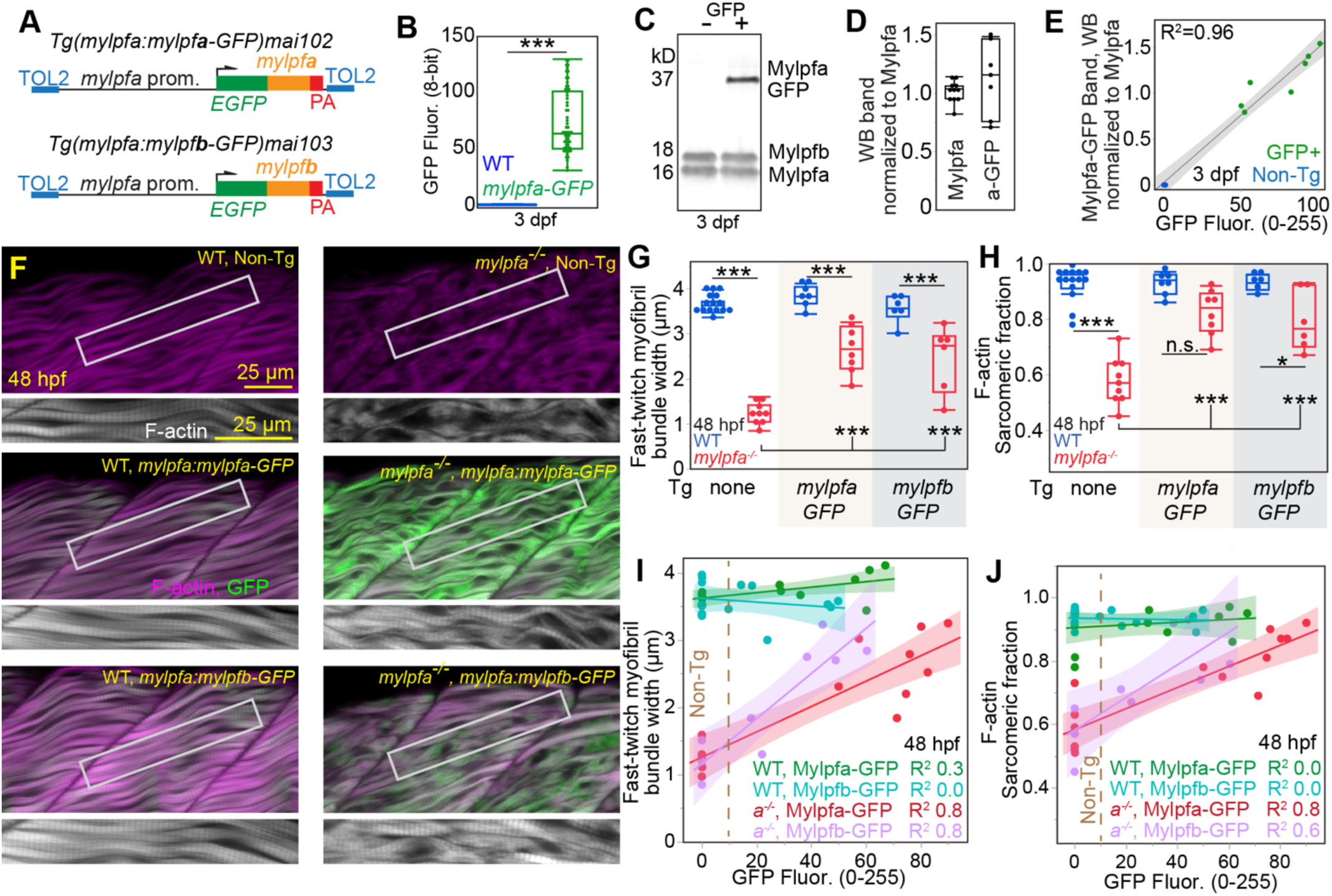
Expression of either *mylpfb-GFP* or *mylpfa-GFP* can rescue *mylpfa^-/-^* myofibrils to a degree proportionate with GFP abundance. **(A)** Schematic illustrating the *Tg*(*mylpfa:mylpfa-GFP)mai102* and *Tg(mylpfa:mylpfb-GFP)mai103* transgenes. **(B)** Box plot showing the variation of fluorescence brightness in the *mylpfa-GFP* transgenic animals and their non-transgenic siblings. The animals imaged for B were pooled into groups of 10 and run on acrylamide gels for western blotting. **(C)** Example of a western blot of Mylpf protein from the wild-type fish and their Mylpfa-GFP+ siblings at 3 dpf shows that the transgene produces one thin band that is shifted upwards by the addition of GFP. **(D)** Plot showing quantification of the Mylpfa, and Mylpfa-GFP bands, normalized to the Mylpfa. **(E)** Correlation plot showing that average fluorescence intensity from these pools corresponds precisely with band intensity in the western blots. **(F)** Confocal slices through fast-twitch muscle with green GFP and magenta phalloidin (F-actin). Zoomed inset shows the F-actin channel at 48 hpf. Panels show representative wild-type and *mylpfa^-/-^*animals that are non-transgenic (top), express the *mylpfa-GFP* transgene (middle), or the *mylpfb-GFP* transgene (bottom). **(G)** Statistical comparisons of myofibril bundle (m.b.) widths in 2 dpf animals. **(H)** Statistical comparisons of the F-actin sarcomeric fraction for these same samples. **(I)** Scatterplot showing correlations between GFP brightness and myofibril bundle width at 2 dpf. **(J)** Scatterplot shows correlations between GFP brightness and the F-actin sarcomeric fraction in the same samples. Points in these panels represent statistical N, which is the individual animal (B, G-J) or pool of animals (C-E). Significance thresholds are determined by Tukey-Kramer HSD comparisons after one-way ANOVA: n.s. P>0.1, * indicates P<0.05, *** P<0.001.

### Mylpf activity is conserved between zebrafish and human

Since Mylpfa and Mylpfb have 80-81% amino acid identity with human MYLPF, we reasoned that Mylpf protein function may be conserved in these two vertebrate species. To test the compatibility of human MYLPF with zebrafish genetics, we generated a transgene that expresses GFP-fused MYLPF under the *mylpfa* promoter, *Tg(mylpfa:MYLPF-GFP)mai104* (Figure 8A). We find that this transgene is capable of restoring myofibril bundle size in the *mylpfa^-/-^* mutant, to a degree that is linearly proportionate with GFP fluorescence, up to the wild-type level (Figure 8B-E). To compare rescue experiments, we scaled each to the non-transgenic wild-type mean for that experiment (2 or 6 dpf). Across timepoints, we find no significant difference in slope between animals expressing MYLPF-GFP, Mylpfa-GFP, and Mylpfb-GFP on scaled myofibril bundle width (Figure 8F). This analysis shows that human MYLPF protein can replace zebrafish Mylpfa and suggests that it acts with an efficiency that is similar to the zebrafish Mylpfa and Mylpfb proteins.

**Figure 8:**
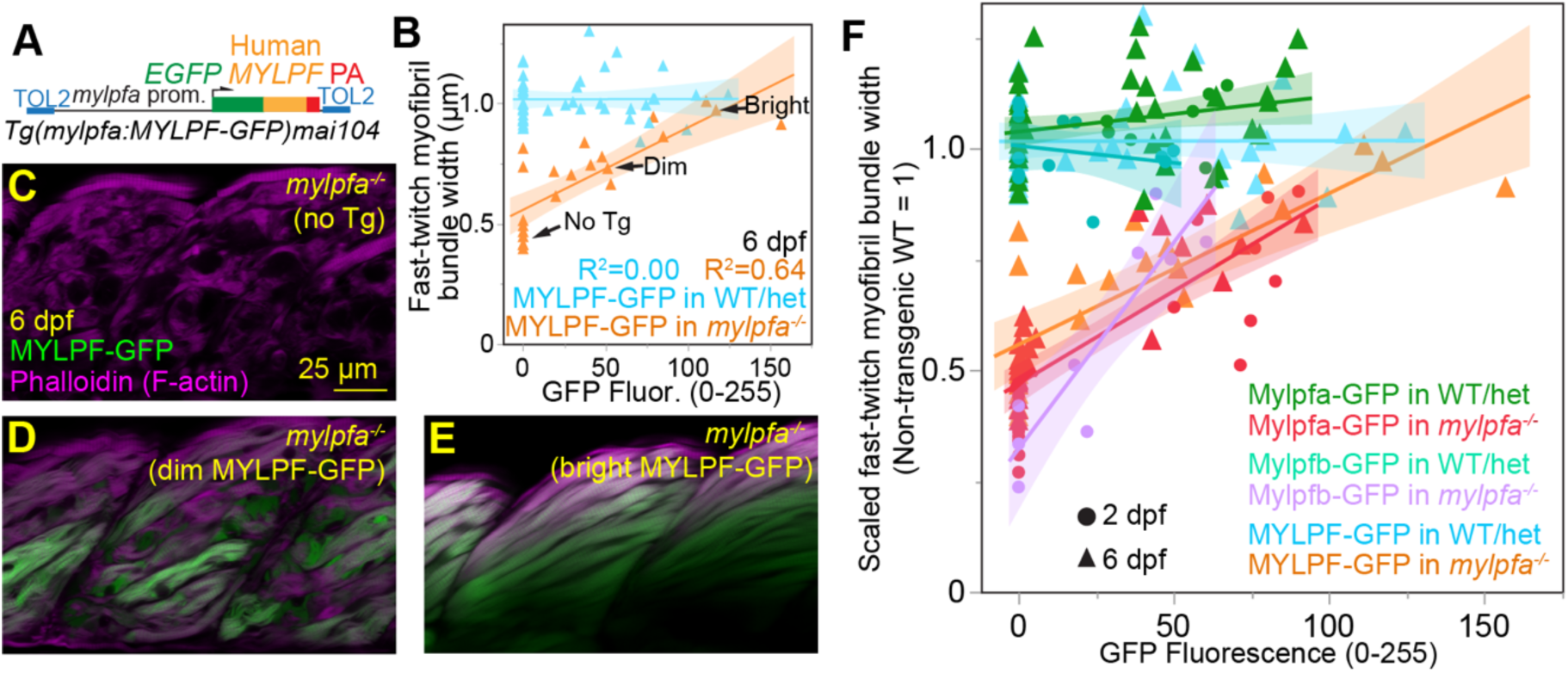
Expression of GFP-fused human MYLPF is sufficient to restore fast-twitch myofibril formation in the *mylpfa^-/-^* mutant. **(A)** Schematic illustrating the *Tg*(*mylpfa:MYLPF-GFP)mai104* transgene. **(B)** Scatterplot showing correlations between GFP brightness and the fast-twitch myofibril bundle width from an intercross of *mylpfa^-/+^* heterozygotes segregating this transgene. **(C-E)** Examples of confocal images of *mylpfa^-/-^*mutant animals that have no GFP expression (C), dim GFP expression (D), and bright GFP expression (E). Quantification of these individuals is shown in B. **(F)** Scatterplot of m.b. width data from all *mylpfa-GFP*, *mylpfb-GFP*, and *MYLPF-GFP* transgenic experiments in this study (Figure S12), normalized to the mean wild-type value per experiment. We find similar slopes in the *mylpfa^-/-^* mutant animals bearing these transgenes, suggesting that these three Mylpf genes act with equivalent efficiency. None of these three transgenic lines significantly influence the widths of myofibrils in wild-type (*mylpfa^+/+^*) animals. Points in panels B and F represent individual animals.

### A MYLPF allele found in DA does not support myofibril formation

We tested whether a DA-causing MYLPF allele can rescue muscle structure as efficiently as its wild-type counterpart. This allele, encoding protein variant G163S, is inherited heterozygously in two individuals with DA, suggesting that partial loss of function may lead to the disease ^33^. Analysis using F0 transient mosaic animals suggests that the G163S MYLPF variant does not promote myofibril bundle formation in the *mylpfa^-/-^* mutant (Figure 9A-E). Likewise, this did not rescue the F-actin sarcomeric fraction, GFP sarcomeric fraction, nor the correlation between actin and MyHC labels (Figures 9F-H, S11). We confirmed the failure to rescue by live imaging the *mylpfa^-/-^*mutant expressing a stable transgenic, *Tg(mylpfa:MYLPF^G163S^-GFP)mai119* alongside the actin transgenic label *TgBAC(six1b:Lifeact-mTurquoise2)mai121* (Figure 9I-I’’). These findings suggest that a nonfunctional MYLPF allele can cause Distal Arthrogryposis when heterozygous.

**Figure 9:**
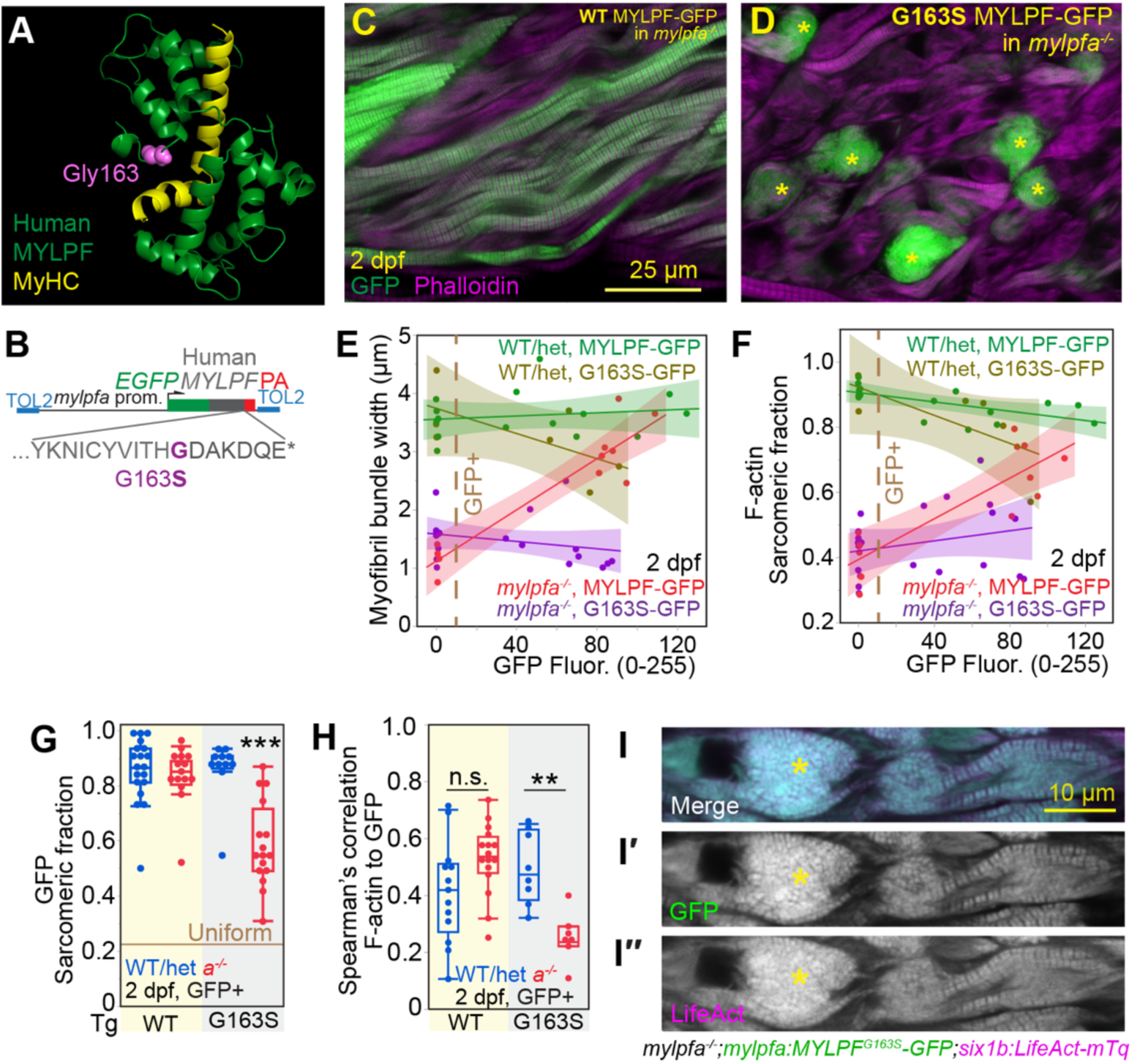
A MYLPF protein variant found in Distal Arthrogryposis fails to rescue the *mylpfa^-/-^* mutant. **(A)** Predicted MYLPF structure, with pink color and filled space highlighting the position of Glycine at residue 163. **(B)** Illustration of a construct that expresses GFP-fused human MYLPF or the G163S variant MYLPF. **(C-D)** Representative images of *mylpfa^-/-^* mutants that were injected with plasmid expressing MYLPF-GFP protein (C) or the G163S variant (D), resulting in mosaic GFP expression. Asterisks indicate balls of highly disordered GFP fluorescence signal, which are only seen in the G163S MYLPF protein variant. **(E)** Scatterplot showing correlates between myofibril width and GFP fluorescence for both injected constructs in the *mylpfa^-/-^*mutant and their WT/het siblings. **(F)** Scatterplot showing the correspondence between F-actin sarcomeric fraction and GFP fluorescence in the *mylpfa^-/-^*mutant and their WT/het siblings. **(G)** Box plots showing the degree to which wild-type MYLPF-GFP or the G163S MYLPF-GFP localizes to a sarcomeric pattern. **(H)** Plot of Spearman’s correlation between GFP and F-actin in these GFP-positive animals. **(I-I’’)** Representative myofibers from a *mylpfa^-/-^*mutant expressing the G163S MYLPF variant from the stable transgenic allele *Tg(mylpfa:MYLPFG163S-GFP)mai119* alongside the live actin marker *TgBAC(six1b:LifeAct-mTq)mai121*. The GFP signal appears mildly sarcomeric but it is not ordered into myofibrils, despite expression at a high level (I’). The LifeAct-mTq signal also appears disordered (I’’). Modest bleed-through from GFP into the mTq channel precludes colocalization analysis. Each point in E-H represents a statistical N, which is the mean of measurements from one animal. Graphs E and F show remeasurements of GFP-negative cells (GFP<10) and GFP-positive cells (GFP>10) within the same plot, divided by a dotted brown line. Significance thresholds in G and H are determined by Tukey-Kramer HSD comparisons after ANOVA: n.s. is P>0.1, ** indicates P<0.01, *** indicates P<0.001. Scalebar in C is for C-D, scalebar in I is for I-I’’.

### Mylpf shows consistent effects on muscle structure from 2 to 6 dpf

We hypothesized that Mylpf dosage may regulate sarcomere assembly with equal potency throughout early development. To understand elements of Mylpf function that may be constant through early development, we scaled our *mylpfa;mylpfb* mutant datasets to the wild-type value at each timepoint and then overlaid measurements, yielding a unified regression model with upper asymptote around 1, EC50 near 0.5, and Hill slope >4 (Figure 10A). We then tested this model by comparing it to findings from animals that express Mylpfa-GFP, Mylpfb-GFP, or MYLPF-GFP at 2 or 6 dpf. To model dosage in individual animals, we converted GFP brightness in each animal to a predicted Mylpf dosage, scaled using the 2:1 ratio of Mylpfa to Mylpfb protein (Figures 10B, S12). The transgenic dataset does not form a lower asymptote because it was generated using the *mylpfa^-/-^* mutant line. We supplied a lower asymptote from the *mylpfa;mylpfb* mutant series, to generate a sigmoidal curve suitable for testing the other three Hill parameters (Figures 10B, S12). The transgenic regression shows an upper asymptote of 1, supporting our claim that Mylpf is insufficient to expand myofibril widths (Figure 10B). In support of our claim that 50% Mylpf dosage impacts phenotype, the transgenic regression shows an EC50 that is close to 0.5. Myofibril bundle widths show a higher EC50 and lower Hs value in transgenic analysis compared to knockout analysis, suggesting that the GFP tag impedes Mylpf efficiency (Figure 10A, B), though we do not detect this difference using the sarcomeric fraction measure (Figure S12). Across all analyses, we find a Hill slope that is considerably larger than one, supporting our claim that Mylpf proteins cooperatively promote the cytoskeletal arrangements that strengthen muscle throughout the first week of development in zebrafish, including myofilament localization, sarcomere formation, myofibril widening, and myofibril bundle growth.

**Figure 10:**
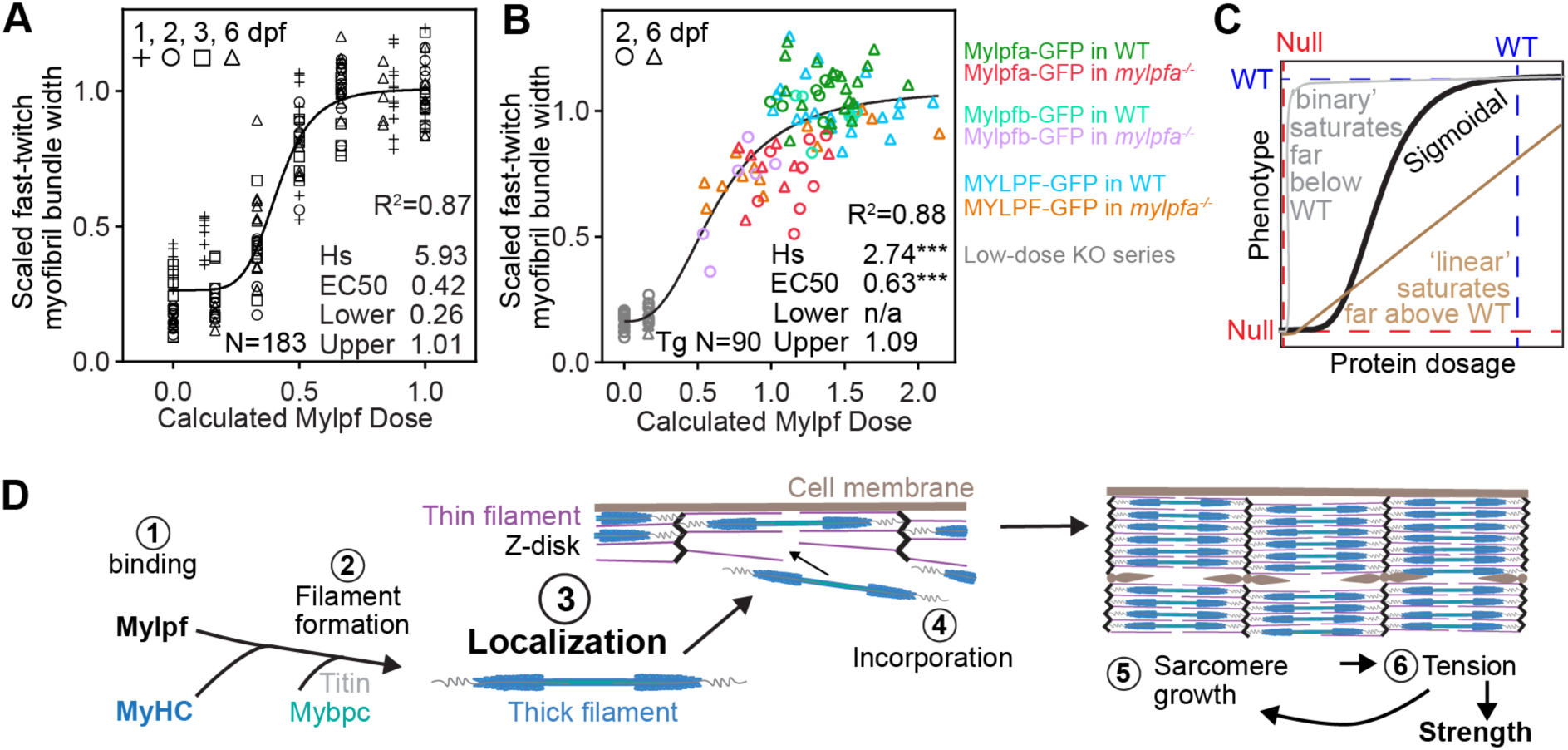
Modeling the impact of Mylpf on muscle structure. Scatterplot graphs, with Y-axis showing the myofibril bundle (m.b.) widths scaled to the non-transgenic wild-type level per experiment, and the X axis showing the Mylpf dosage as calculated using formulas shown in Figure S12. **(A)** A plot of data from the *mylpfa;mylpfb* mutant series shows strong overlap at each timepoint examined (1, 2, 3, and 6 dpf) using datapoints presented in Figure 5B-D. The Hill regression model shows a high correlate across timepoints. **(B)** A plot of the transgenic data, excluding GFP <10, alongside the most severe Mylpf loss of function alleles from the mutant lines (*mylpfa^-/-^;mylpfb^+/-^* and *mylpfa^-/-^;mylpfb^-/-^*, gray dots). This dataset also produces a Hill regression with a high R^2^, high Hill slope and moderate EC50. Asterisks on Hs and EC50 values reflect fit of the dataset in panel B for transgenic animals to the regression curve calculated in A for the *mylpfa;mylpfb* mutant series; significance threshold. Asterisks, *** indicates P<0.001. **(C)** Examples of three potential types of phenotypic responses to protein dose. Mylpf appears to have a sigmoidal response curve with a steep slope at inflection and an EC50 value somewhat below 50% protein dosage. **(D)** A model for how Mylpf may cooperatively promote sarcomere and myofibril bundle assembly. Our findings support a role for Mylpf in MyHC localization and Mylpf may cooperatively promote other steps in this process.

## Discussion

Our central finding is that Mylpf function is essential for the defining feature of striated muscle, the overlap of sliding filaments ^37,38^. We show that the protein is necessary but not sufficient for myofibril formation in fast-twitch muscle; however, this binary language does not fully capture the sensitivity of muscle development to Mylpf loss. Our work was initiated by the intriguing finding that *mylpfa* is required for muscle structure, but *mylpfb* seems not to be. This difference could be explained by the timing of gene expression, since *mylpfa* mRNA abundance climbs faster than *mylpfb,* with a 7:1 ratio at 1 dpf and 2:1 ratio by 2 dpf. However, the large discrepancy at 1 dpf does not explain why the phenotypic differences between *mylpfa^-/-^* and *mylpfb^-/-^*persist so long after *mylpfb* levels increase. Another simple explanation is that Mylpfa protein has some function that Mylpfb is lacking; however, that model would not explain our observation that both Mylpfa-GFP and Mylpfb-GFP rescue the *mylpfa^-/-^* mutant with similar efficiency. Instead, we propose that Mylpf protein dosage can explain the phenotypes we observed across all Mylpf genotypes and timepoints, when modeled using a Hill regression (Figure 10A, B). This regression model is a popular tool for biochemical analysis and has also been used previously to investigate the dose sensitivity of gene promoters *in vivo* ^1,39^. This model suggests that the *mylpfb^-/-^* mutant expresses total Mylpf protein just slightly above its functional saturation point, around two thirds the wild-type level. For Mylpf, we find an overtly sigmoidal relationship between dosage and phenotype from the null to wild-type genotypes; however, some Hill models can seem nearly binary or nearly linear in the range between null and wild-type (Figure 10C). An overtly sigmoidal relationship could arise because Mylpf is a structural protein that functions stoichiometrically in proportion to other sarcomeric proteins, consistent with recent proteomic work in cardiac muscle ^40^. These findings raise the intriguing possibility that many other muscle proteins have stoichiometric dose-dependency, because they function together across entire myofibril bundles during sarcomere formation and force production.

The relationship between Mylpf protein dosage and sarcomeric phenotype has a large Hill slope, suggesting that Mylpf proteins function cooperatively during sarcomere formation ^41^. The Hill equation is traditionally used to describe cooperativity in ligand-substrate binding, though later steps may also be synergistic (Figure 10D). For instance, each Mylpf protein should help to stabilize MyHC during thick filament production (step 2). We think that the core step regulated by Mylpf may be myofilament localization (step 3), since we see poor colocalization of F-actin with MyHC across multiple timepoints of development. Alternatively, the myofilaments may be returned to the cytoplasm after briefly localizing. Both scenarios result in mislocalized thick filaments, which cannot interdigitate with thin filaments (step 4) ^4^, leading to loss of the tension-based feedback loops shown to regulate sarcomere growth (steps 5, 6) ^6,42^. These feedback loops may be disrupted by impaired Mylpf stability, since point mutations in RLCs are known to affect step size ^14^. Prior work indicated that Mylpf also controls overall muscle growth, since the mouse *Mylpf^-/-^*knockout is born without skeletal muscle ^22^ and myosin heavy chains are needed to support myofibril organization and muscle growth ^43–45^. The *mylpfa^-/-^* mutant shows reduction in somite growth by 6 dpf, when assessed in live animals. We find similar average measurements using phalloidin on fixed samples, though here the difference in size between genotypes does not rise to statistical significance. The *mylpfa^-/-^*muscle growth defect is consistent with prior work showing that MyHC itself is required for myofiber growth and the known feedback loops between strength and growth ^43,44^. Furthermore, the effects on myofibril bundle growth are consistently larger than those on somite growth. We think that the primary role for Mylpf is in sarcomere formation, because this function occurs immediately and has cascading effects.

Given its complete sarcomere loss in fast-twitch muscle (over 95% of total muscle), we were surprised to find the *mylpfa^-/-^;mylpfb^-/-^*double mutant moving as far as its wild-type siblings. The *mylpfa^-/-^*mutant and *mylpfa^-/-^;mylpfb^-/-^* double mutant have classic signs of fast-twitch deficiency including an absence of C-bends and impaired escape response, consistent with previous work on zebrafish showing loss of function in fast-twitch muscle ^33,46,47^. We propose that the *mylpfa^-/-^;mylpfb^-/-^* double mutant swims so far because they increase the frequency of slow-twitch movements, which arise independently of fast-twitch activity ^48^. Some genotypes, such as the *mylpfa^+/-^;mylpfb^+/-^*double heterozygote upregulate these slower movements and have functional fast-twitch muscle, allowing them to outpace their wild-type siblings. Consistent with our model, zebrafish lacking fast-twitch acetylcholine receptors upregulate slow-twitch synapses and behaviors by adult stages ^46^. Alternatively, the behavioral compensation in Mylpf-deficient animals may be caused by excess exercise of the slow-twitch muscle fibers ^28^. The importance of muscle hypertrophy and neural activity could be resolved by investigating muscle strength, calcium signaling, and synapse morphology through time in Mylpf mutants. Investigating this compensatory mechanism may provide crucial insights for human health, since many muscle diseases disproportionately affect the fast-twitch fiber type^28^.

The impact of Mylpf dosage on muscle formation may not be restricted to zebrafish. Our G163S findings suggest that the allele has little or no activity, so a person heterozygous for this variant may have dominantly inherited DA because they have around half the normal MYLPF activity. Likewise, we show that zebrafish with 50% Mylpf dose (the *mylpfa^+/-^;mylpfb^+/-^*double heterozygote) have impaired myofibril bundle assembly. Similarly, a heterozygous effect is also found in *Drosophila,* wherein RLC^+/-^ flight muscles (fast-twitch) show impaired myofibril assembly ^17^. No defects were reported in a study of the mouse *Mylpf^+/-^* heterozygote, although embryonic myofibril bundle widths were not measured ^22^. However, dosage appears to be broadly relevant in mammals, since studies of pigs and buffalo show that Mylpf mRNA levels can predict meat quality ^49,50^. Myosin light chain dosage also appears to impact heart muscles, since activation of cardiac myosin light chain kinase is necessary and sufficient to initiate the formation of sarcomeres in cultured heart cells ^51^. We propose that the relationship between sarcomeric protein dosage and myofibril expansion may have applications in many contexts where muscle strength is desired including farming, exercise physiology, and human health.

## Supporting information

VIdeo 1: Confocal stacks showing differences in muscle structure between wild-type and mylpfa-/- sibling animals.

Video 2: The mylpfa-/- mutant shows an impaired escape response.

## Acknowledgements

The fish facilities at the University of Maine and The Ohio State University (OSU) provided excellent animal care. Dr. Joy-El Talbot from Iris Data Solutions developed a FIJI macro for sarcomere fraction quantification. Dr. April DeLaurier provided feedback in the writing process. Dr. Clarissa Henry generously provided mentorship and resources to the Talbot lab. Dr. Thomas Gallagher and Collin Whitlock facilitated data sharing between OSU and the University of Maine. Gursimran Dhillon provided training on technique and data analysis for RT-qPCR. Paula Monsma and Dr. Sarah Cole supported imaging on the electron and confocal microscopes at OSU. Dr. Zhengxin Ma supported training and usage of the Zeiss LSM980 confocal in the Microscopy and Image Analysis Core facility at the University of Maine (RRID:SCR_025784). Antibodies mMac ^52^ and A4.1025 ^53^ were provided by DSHB at the University of Iowa. This project was supported by a University of Maine Institute of Medicine Seed Grant, and NIH grant R15AR081019 to JCT, NIH R15GM140409 to JBK, by NIH grants GM088041 and GM117964 to SLA, and COBRE 1P20GM144265 to JBK and JCT.

## Materials and Methods

### Zebrafish husbandry and established strains

Fish strains were maintained using standard methods ^54^. All animal protocols used in this study were approved by the Institutional Animal Care and Use Committees at The Ohio State University (2012A00000113) and the University of Maine (A2022-09-05, A2025-10-01). Zebrafish staging followed established metrics ^55^. We genotyped the *mylpfa^oz^*^30^ 20 bp deletion allele ^33^ using primers (F 5’-TCTCTACAGGCCAGCTGAATG’3’, R 5’-ACCCTTCAACTTCTCTCCGAAC’3’) that amplify a 116 bp product in wild-type fish and a 96 bp product in the *mylpfa^oz30^* homozygote. We genotyped the *mylpfa^oz43^* +1 bp insertion allele ^33^, (primers 5’-GCTTCATTGCTGTCAGGATAGAG-3’ and 5’-ACCCTTCAACTTCTCTCCGAAC-3’), followed by digestion with BsmFI restriction enzyme (NEB, New England Biolabs) which cuts the wild-type but not mutant amplicon. Homozygotes for both *mylpfa^oz30^*and *mylpfa^oz43^* can be consistently sorted from wild-type siblings based on pectoral fin immobility, a slowed escape response, and reduced birefringence ^33^. The *mylpfa^oz30^*allele was used for all *mylpfa^-/-^* mutant analysis in this study, except for Figure S5E. Zebrafish lines in this study were maintained on the AB wild-type background.

### Transgene construction

The four Mylpf-containing plasmids constructed for mosaic analysis in this study were *pMylpfa:mylpfa-GFP*, *pMylpfa:mylpfb-GFP*, *pMylpfa:WT-MYLPF-GFP*, and *pMylpfa:G163S-MYLPF-GFP.* Briefly, we obtained GeneArt-strings (Invitrogen) of the sequence encoding Mylpfa, Mylpfb, wild-type human MYLPF or MYLPF p.Gly163Ser (G163S), proteins. These coding sequences were inserted into a plasmid containing the *mylpfa* promoter flanked by TOL2 sites ^34,56,57^; GFP was linked to the 5’ end of Mylpf genes, connected via a sequence encoding a short flexible peptide (GGGGSGAT). The LifeAct transgene was made by modifying a previously described six1b BAC ^58^ by integrating coding sequence for LifeAct-mTurquoise2 (LifeAct-mTq) protein after the *six1b* start codon. Plasmid and BAC injection was accompanied by co-injection of zebrafish-optimized nuclear transposase ZT2TP ^59,60^. For transient analysis, animals with high levels of mosaicism were selected for imaging and all experiments were repeated on at least two independent injection days. Stable transgenic lines were generated for five transgenes, *Tg*(*mylpfa:mylpfa-GFP*)*mai102*, *Tg*(*mylpfa:mylpfb-GFP*)*mai103, Tg(mylpfa:MYLPF-GFP)mai104, Tg(mylpfa:MYLPF^G163S^-GFP)mai119,* and *TgBAC(six1b:Lifeact-mTurquoise2)mai121*. For stable transgenic analysis, progeny from one founder (*Lifeact-mTurquoise2*), two founders (*mylpfa-GFP, MYLPF-GFP*) or four founders (*mylpfb-GFP, MYLPF^G163S^-GFP*) were analyzed.

### Construction of *mylpfb^oz39^*

The *mylpfb^oz39^* mutation was generated using established CRISPR-Cas9 protocols ^61^. One-cell embryos were co-injected with Cas9 mRNA and guide RNA targeting sequences in exon 2 (5’-GGAAAACAGTAAAGTTGATG –3’), raised to adulthood, and outcrossed to identify germline-transmitting founders. F1 progeny were screened using High-Resolution Melting Analysis (HRMA) (primers 5’-CCCTCTCTAAAACAAACAGGCTTTC-3’ and 5’-GGTAAGTGAAGATTTGGACAACTC-3’) to identify founders carrying the frameshifting –5 bp deletion *mylpfb^oz39^* allele. The allele sequence was confirmed via Sanger sequencing of homozygotes and the founder was outcrossed for at least two generations before conducting experiments. We identify the *mylpfb^oz39^*lesion via TaqMan real-time PCR genotyping assays. The TaqMan real-time PCR method employs custom primers and TaqMan probes to distinguish between wild-type and *mylpfb* mutant amplicons. The forward and reverse primers used are 5’-AAGGATGATTTGAGGGATGTG-3’ and 5’-TTTCACCAAACATGGTAAGGAA-3’, respectively; fluorogenic probes are (56-FAM/CC +CA+T +C+AA +CT+T TAC /3IABkFQ/) for wild-type amplicons and (5SUN/CC C+CA +T+T+T A+CT G+TT /3IABkFQ/) for mutant.

### RNA isolation and cDNA synthesis

We determined absolute mRNA expression levels across developmental stages (24–72 hours post-fertilization, hpf) using total RNA isolated from AB wild-type embryos. Embryo pools of 40 (24 hpf), 30 (36 hpf), 20 (48 hpf), and 10 (72 hpf) were homogenized in Trizol using 0.5 mm zirconium beads (Laboratory Supply Network), and RNA was subsequently purified using the Direct-zol RNA Miniprep Kit (Zymo Research). We assessed the relative mRNA expression in the *mylpfa;mylpfb* mutant series using decapitated 48 hpf embryos obtained from incross of adult *mylpfa^+/-^;mylpfb^+/-^*double heterozygotes. The ‘head’ DNA was used for genotyping, while bodies were immediately snap-frozen at −80°C in individual tubes. After genotyping, RNA was extracted from pools of three embryos per genotype. Purified total RNA (1 µg for AB wild-type or 100 ng of the *mylpfa;mylpfb* mutant series) was reverse transcribed using the SuperScript IV Reverse Transcriptase Kit (Invitrogen), along with oligodT_20_ primers (Invitrogen) and murine RNase inhibitor (NEB), according to the manufacturer’s protocol.

### RT-qPCR

RT-qPCR was performed using SYBR Green master mix (Bio-Rad) on CFX384 thermocycler (Bio-Rad), with 60°C annealing and 30 second extension. Each reaction used 15 ng of cDNA in an 10 μl volume, performed in technical triplicate. The *mylpfa* qPCR generates an 80 bp amplicon using the Forward primer, 5’-TCAATGGGCCAGCTGAATGT –3′ and Reverse primer, 5′ – ACGGTGAAGTTGATTGGGCC –3′. The *mylpfb* qPCR generates an 83 bp amplicon using the Forward primer, 5’-AGGAATACAAGGAGGCTTTCACA –3′ and Reverse primer, 5′ – TGCCAGCACATCCCTCAAAT – 3′. The *rpl13a* transcript was selected as the reference gene due to its stable expression during zebrafish embryonic development ^62^. The *rpl13a* qPCR uses primers: Forward 5’-TCTGGAGGACTGTAAGAGGTATGC-3′ and Reverse 5’-AGACGCACAATCTTGAGAGCAG-3′. Relative mRNA quantification was performed using the 2^-ΔΔCq^ method ^63^ after normalization to the mean of all wild-type groups for each target gene. Each experiment was repeated on at least three biological replicates before statistical analysis.

### Absolute quantification of *mylpfa* and *mylpfb* mRNA

For absolute quantification of mRNA copy number, we generated *mylpfa* and *mylpfb* DNA templates (called “standard DNA”) from 72 hpf cDNA. For *mylpfa* standard DNA, a 563 bp was amplified from the zebrafish *mylpfa* gene using the Forward primer, 5’-ACTACGCGGCTTCAGACTTC –3′ and Reverse primer, 5′ – TCCTTCTCCTCTCCGTGTGT – 3′. For *mylpfb* standard DNA, a 555 bp was amplified from the zebrafish *mylpfb* gene using the Forward primer, 5’-TGTCAGTCACATCAACGTGGA –3′ and Reverse primer, 5′ – CCTCTCCATGTGTGATGACGT – 3′. DNA standards were purified using a gel extraction kit (Thermo Fisher) and sequence was confirmed. To calibrate for copy number, we amplified a series of standard DNA concentrations spread across ten four-fold dilutions on each qPCR plate, with predicted concentration ranging from 4.5 copies/reaction to 4.92×10⁶ copies/reaction. The graph of the curve was used to calculate the number of cDNA molecules tested in the same reaction plate as the standard, and a ratio between *mylpfa* and *mylpfb* copies is presented.

### Immunohistochemistry and HCR ISH

Whole mount immunohistochemistry used established markers and techniques ^64^. Briefly, the larvae were fixed in 4% paraformaldehyde and washed in PBST (1X PBS, 0.1% Tween-20) before immunolabel. Embryos older than 1 dpf were gently permeabilized using a brief treatment with 0.001% Proteinase-K, washed repeatedly in PBST, then blocked for at least two hours before overnight incubation at 4°C with primary antibodies diluted in blocking solution. We used primary antibodies for the myonuclei (1:500, Rbfox1l) ^65^; α-Actinin (1:500, A7732, Sigma); Myomesin [1:30, mMac, Developmental Studies Hybridoma Bank (DSHB)] ^52^, MyHC (1:1000, A4.1025, DSHB)^53^. F-actin labeling uses Alexa Fluor-568 conjugated phalloidin, added during the secondary antibody step (1:50, Thermo Fisher). HCR ISH procedures use established protocols with probes and amplifiers supplied by Molecular Instruments ^66^. Our HCR mixtures included nine unique *mylpfa* probes, five *mylpfb* probes, and twenty *myl10* probes. To compare HCR ISH signal intensity, separate fish were labeled with *mylpfa* and *mylpfb* probes, both in the 488 nm excitation channel, and then imaged with matched settings on a dissecting microscope before analysis in FIJI.

### Protein modeling

Mylpf proteins were modeled in complex with the binding region of zebrafish Myhz1.3 (zebrafish) or MYH3 (human) using Robetta ^67^ and the models were visualized using PyMol (Schrödinger).

### Antibody production, validation on western blot

The zebrafish Mylpf antibody was generated by Boster Bio, using zebrafish Mylpfa (AA 14 to 167 of NP_571263.1) in complex with a Mylpf-binding region that is found in several MyHC proteins (RRESIYTIQYNIRSFMNVKHWPWMKVYYKIKPL). Western blot in the Talbot Lab showed only one band each for Mylpfa (16 kD) and Mylpfb (18 kD) and no label in the MyHC size (252 kD). For western blotting, animals were raised to the desired stage, then pools or single animal were homogenized in 100 µl (pools) or 20 µl (single) 1X LDS sample buffer solution (Bolt; Invitrogen) using 0.5 mm zirconium beads (Laboratory Supply Network). In the western blot showing Mylpfb and Mylpfa protein abundance in wildtype (AB) animals, we used pools of 40, 30, 20 and 10 animals at 24, 36, 48, and 72 hpf respectively to normalize for zebrafish growth. Prior to western blot, animals were decapitated and deyolked to enrich for somite muscle and provide genotyping materials. For the *mylpfa;mylpfb* mutant series, we ran the protein from one animal per genotype across the nine genotypes generated by incross of *mylpfa^+/-^;mylpfb^+/-^* double heterozygotes. Homogenized samples (20 µl) were run on a 4-20% gradient acrylamide gel (Bio-Rad). The protein samples were transferred to Polyvinylidene difluoride (PVDF) membranes (Millipore Sigma) which were blocked (5% nonfat dry milk), then replaced with primary antibody solution and then secondary antibody solution. Primary antibodies included mouse polyclonal anti-MyHC (A4.1025, DSHB, 1:1000) to detect MyHC protein while rabbit polyclonal anti-Mylpf (Boster, 1:1000) detected both Mylpfa and Mylpfb protein. The secondary antibodies used were donkey-anti-mouse-800 (1:10000 dilution) and donkey-anti-rabbit-680 (1:10000 dilution). The resulting fluorescence was measured by scanning the blot with an Odyssey Infrared Imager (LICOR). The Mylpfa to Mylpfb ratio was quantified using GelBox software ^68^ and FIJI ^69^.

### Confocal imaging

Confocal images in Figure S5 were collected using an inverted Nikon TiE microscope equipped with an Andor Revolution WD spinning disk confocal system. Confocal images in Figure 2A-D’ and S8 were collected using a Zeiss LSM980 confocal microscope. All other confocal images were acquired using a Leica TCS Sp8 confocal microscope and processed with Lightning in LasX software. Myotomes were imaged over the mid yolk tube region of embryos and larvae. Image settings including laser power, gain, and confocal export protocols were standardized for each experiment. The channels tool is used to generate colorblind friendly images, such as converting red to magenta.

### GFP brightness analysis

We evaluated the GFP brightness using the mean grayscale value of the GFP channel in FIJI. For stable transgenic lines, we drew ROI around the fast-twitch region of the somite. In mosaic animals, we drew ROI around each muscle fiber because there was often a high level of variation within a single image. Muscle was considered non-transgenic if the GFP brightness is <10 (0-255) and considered transgenic when in the 10-255 range. MATLAB was used to calculate the average of each mean grayscale value per image followed by statistical comparisons in JMP software.

### Sarcomeric fraction calculation

We developed a metric to quantify the degree to which muscle proteins localize to sarcomeres, termed the sarcomeric fraction (Figure S4). Grayscale intensities were calculated across Region of Interest (ROI) lines along the length of a myofiber in FIJI, with ten to thirty ROIs measured per image. The periodicity of these intensities was determined using the ‘distance between peaks’ function in MATLAB. Peak-to-peak distances per intensity were determined per image, and bootstrap confidence intervals were calculated using variation between images, shown via histogram to represent the Fraction of Peaks/micron (µm). Signal was considered ‘sarcomeric’ in different intervals (’bin sets’) for markers with full-sarcomere (F-actin, Myomesin, Actinin) or half-sarcomere repeat (MyHC, Mylpf-GFP), or an even blend of the two patterns (LifeAct) (Figure S3). Sarcomeric LifeAct shows peak signals in the 0.7-1.1 µm and 1.55-2.30 µm range. Sarcomeric phalloidin (F-actin) also labels Z-disks and thin filaments, showing peak signals in the 0.7-1.0 µm range and 1.55-2.15 µm range. For consistency, we used the F-actin bin set for Myomesin and Actinin (Figure S4). Sarcomeric MyHC and Mylpf-GFP show two peak signals in bins 0.6-1.2 µm and 1.75-2.05 µm (Figure S4). These bins were determined by examining histogram plots of the localization of sarcomeric signals during preliminary analysis (examples in Figure S4) and then applied evenly to subsequent analyses. The sarcomeric bins occupy 0.9 µm of the 4 µm sampled, so a uniformly distributed label would score 0.225 (shown with a brown line in graphs) whereas a 1.0 indicates full localization of signal to sarcomeres. The sarcomeric fraction per animal represents the ratio of image periodicity within the sarcomeric bin to the overall periodicity of that marker, averaged across all ROIs (Figure S4).

### Colocalization

The extent of colocalization between confocal channels was determined using the COLOC 2 tool in FIJI. Briefly, we identified large regions of interest (ROI) areas where labeling for the markers compared was homogenously bright. This selection avoided penetration issues, because the MyHC label is brighter at the surface of somites than it is deeper. Typically, the area fills at least half of the fast-twitch muscle region of a somite. We recorded the Spearman’s rank correlation coefficient, Mander’s tM1, and Mander’s tM2 for all images. Measurement was repeated if a large disconnect was found between tM1 and tM2 suggesting that an ROI extended to a region of poor labeling; slight ROI adjustments brought the Mander’s measures into alignment and produced a trustworthy Spearman’s correlation. These Spearman’s correlations were tabulated and subjected to our standard statistical analysis.

### Cross-sectional analysis

To analyze muscle size in cross-sectional view, confocal stacks were acquired with 0.5 µm step size, then orthogonal views were exported to FIJI. Somite area is the total area enclosed within ROIs drawn around the somite. Based on cell position and markers we can also identify fiber types, with the superficial layer being slow-twitch and all deeper layers being fast-twitch ^26^.To measure the cross-sectional area of myofibrils within slow-twitch fibers, we drew ROI encircling the desired fiber type and then performed auto local thresholding with the Bernsend method, setting radius to 15 and the T parameter to 7.5, before measuring area within threshold. To determine protein positions within fast-twitch muscle fibers (Figure 4G-J), a mask was drawn manually, circling the borders of each cell. A custom MATLAB script determined image brightness at each pixel, moving from the periphery of the cells to the center. These edge-center distances were scaled to micron lengths, averaged, and bootstrap confidence intervals were calculated across samples of a given genotype. Graphs with confidence intervals were then overlaid for each genotype.

### Myofibril bundle width measurements

To measure myofibril width, we drew ROI lines across the narrow axis of the myofibril in FIJI. We typically measured thirty myofibrils bundles per image. Each measurement was made at a widened point of each myofibril, with measurements distributed evenly across the fast-twitch muscle region. For mosaic animals we measured both GFP+ and GFP-negative myofibril bundles within the same confocal slice. Images were included only if they contained at least ten GFP+ myofibril bundles. For stable transgenic lines, we compared measurements from non-transgenic and transgenic siblings. We generated a custom FIJI macro to simplify measurement and compile all measures from a given image. These measurements were averaged per image using MATLAB and then imported to JMP software for statistical analysis.

### Transmission electron microscopy

For TEM, embryos were raised to 72 hpf, fixed in phosphate-buffered glutaraldehyde, resin embedded, thin sectioned, and contrasted using uranyl acetate and Reynold’s lead citrate before imaging an initial dataset on a Tecnai 30 G2 TWIN microscope at the Ohio State University and later dataset on a Tecnai T12 BioTwin microscope at the University of Maine. The two datasets were then merged. In total, we imaged 6 wild-type animals, 6 *mylpfa^-/-^*, 3 *mylpfb^-/-^*, and 6 *mylpfa^-/-^;mylpfb^-/-^* double mutant animals, each with multiple sections. We set the mean of measurements within an animal as the single N for statistical comparisons. Myofibril widths were measured between triads and myofibril bundle widths were measured as in light microscopy; both were measured on low-magnification TEM images to sample variation across several myofibers.

### Behavioral analysis

For tail curvature assessment, individual larvae were placed in a petri dish and prodded with a piece of fishing line, imaged using a GoPro Hero camera with a mounting ring (H12Pro, Back-Bone) set to 240 frames/sec. The frame with highest curvature was transferred to FIJI and curvature was measured along the length of the trunk using the Kappa plugin ^70^. For speed analysis, larvae were placed one per well into a 12-well plate with 3 ml of facility water at 6 dpf. Then, they were imaged in a DanioVision system (Noldus) equipped with a magnifier for improved resolution. The plates were incubated at 28°C in the preheated chamber. The larvae were imaged using the infrared wavelength every 0.033 seconds for a total of 25 minutes, cycling lights off and on every five minutes. Larvae were tracked using EthoVision XT Version 15.0 video tracking software by Noldus. ‘Distance traveled’ is the sum of distances in this 25-minute time interval. It is important to use software that includes low-speed movement, such as Ethovision. Analysis was limited to speeds under 200 mm/sec because video examination shows that higher-speed measurements are indications of tracking glitches. The ‘highest speed’ was calculated by sorting data numerically and recording the highest number beneath 200 mm/sec. Average speed was the average of all momentary speeds. Speed is calculated per 0.033 second time interval. Movement is summarized using the Microsoft Excel Histogram function, as the percentage of time that fish swim in a wide range of speed ‘bins’. The first bin is set to 1 mm/sec, then 1.09 mm/sec, then bins continue using the formula Y=X+Ln(X), where Y is the current bin value and X is the previous bin value. The larvae swim exponentially less frequently at high speeds than low speeds, so this natural-log based formula allows comparable statistical sampling of the whole speed range (Figure S7). Shown plots integrate data collected from the *mylpfb^+/-^* heterozygous incross, *mylpfa^+/-^;mylpfb^+/-^*double heterozygous incross, and outcrosses of *mylpfa^+/-^;mylpfb^+/-^* to AB (Figure 6D-G) or from *mylpfa^+/-^* incrosses (Figure 6H-J). Each set of crosses was replicated at least three times with consistent results across experiments.

### Statistical analysis

We begin multiple comparisons using ANOVA and then use Tukey-Kramer’s post-hoc comparison of levels. Pairwise comparisons use a two-sided Student’s T-test and matching results are found with the non-parametric Kruskal-Wallis exact test. In all figures, * P<0.05, ** P<0.01, *** P<0.001. Not significant (n.s.) is P>0.1 or P>0.05 as defined in legend. For box plots, the central line shows the median, the upper bound shows the upper quartile, the lower bound shows the lower quartile, and whiskers show 1.5x the interquartile range. Each N represents a different animal or in some cases a pool of animals and are listed explicitly in the source data file. For mosaic animals, the N represents the number of animals scored for GFP+ or GFP-negative cells, respectively. Repeated measures of the same animal are averaged and included as a single N. Statistical comparisons were made in Jmp Pro, GraphPad, or MATLAB software. Logistic regression models used the four-parameter Hill equation in GraphPad with formula y=L+((U-L)*X^Hs)/(X^Hc+EC50^Hs). Variables are defined as phenotypic response (y), dosage (x), upper asymptote (U), lower asymptote (L), EC50, and Hill slope (Hs). Dose modeling in transgenic animals uses the formula WT=1+0.667*(GFP/57), heterozygote=0.667+0.667*(GFP/57), and *mylpfa^-/-^* =0.333+0.667*(GFP/57), with 57 being the average fluorescence in our images.

### Data availability

The data leading to conclusions in this paper are shown in the main text figures and supplementary figures, with source data provided. We are happy to accommodate requests for additional source files.

### Code availability

Code used for data analysis in this manuscript is available upon request and from our GitHub account, https://github.com/MuscleZebrafish/MyofibrilQuant.

