## Supplemental figures and legends for "Myosin light chain proteins cooperatively promote sarcomere growth in fast-twitch muscle"

### Supplemental Figures with legends

**Figure S1:** Although two transcripts are annotated for *mylpfa* and *mylpfb*, only one is detected in embryonic stages. **(A)** Genomic structure of *mylpfa* and *mylpfb*, showing the two transcripts annotated in Ensembl and the location of lesions used in this study. **(B)** Putative mRNA sequences produced by these genes, also showing the locations of frameshifting alleles in *mylpfa* and *mylpfb*. Regions shared between the two potential transcripts are shown, and unique regions are annotated "alt" (red). **(C-F)** RT-PCR of the four potential transcripts spanning regions marked blue in B, on cDNA from wild-type animals at 24, 36, 48, and 72 hpf. Each panel uses a different primer pair that uniquely amplifies a given transcript. We consistently amplify product from the *mylpfa-201* (C) and *mylpfb-201* (D) transcripts. We do not detect any *mylpfa-202* (E), nor *mylpfb-202* (F) transcript at any of the stages examined.

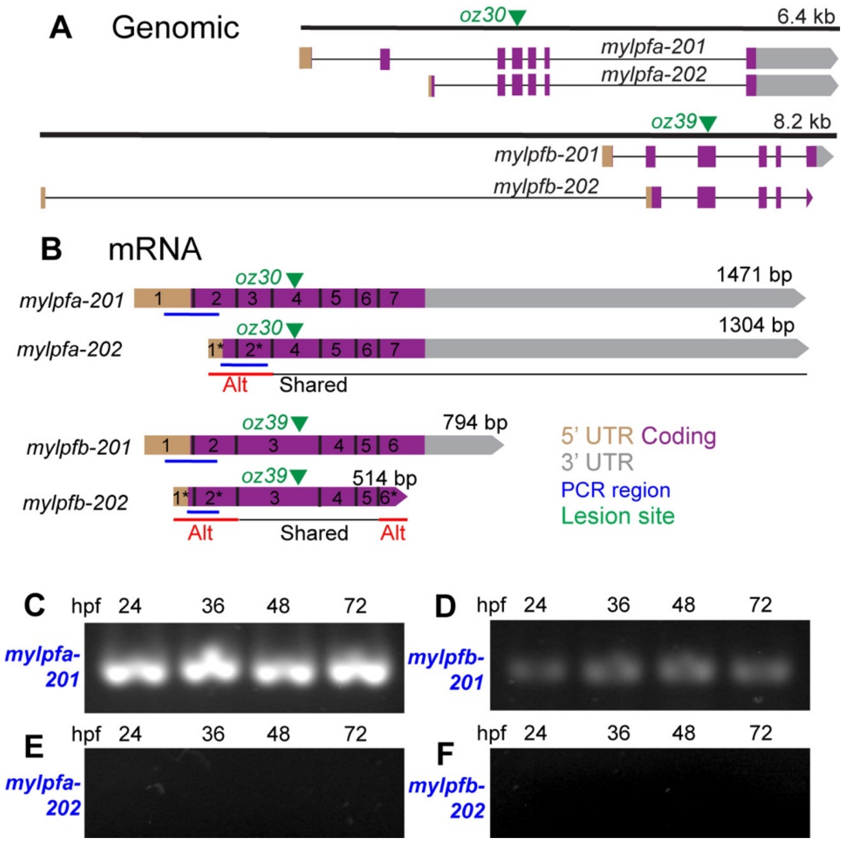

**Figure S2:** Time-course of *mylpfa* and *mylpfb* expression in fast-twitch muscle fibers. Images of HCR ISH for *mylpfa*, *mylpfb*, and the slow muscle marker *myl10.* **(A-C)** Low magnification images highlight that all three genes are expressed specifically in muscle and not in other body locations like the head. **(D-F″)** Confocal slices at higher magnification show that the two Mylpf genes are consistently excluded from slow-twitch fibers, but overlap in all fast-twitch fibers. Both genes are visible in the brightened single-channel inset at the stages examined (D′-F″). The ratio of *mylpfa* to *mylpfb* is somewhat lower in these co-labeled images (detected in 488 and 568 excitation) than in the single-channel ones shown in Figure 1G, H (detected in 488 channel), because of differential sensitivities between 488 and 568 detectors. **(G-G″)** Z-slice from a 20 hpf embryo shown from a transverse view, as a 3-color merge (G), merge of *myl10* (blue) and *mylpfa* (gray) (G**″**), and as a merge of *myl10* (blue) and *mylpfb* (gray) channels (G″). Both genes are present in fast-twitch myofibers, which are medial to slow-twitch fibers by 20 hpf. Scalebar in A is for A-C, D is for D-F, D′ is for D′-F″, and G is for G-G″. Image settings are matched in A-C, D-F, and D′-F′ to allow comparisons between timepoints.

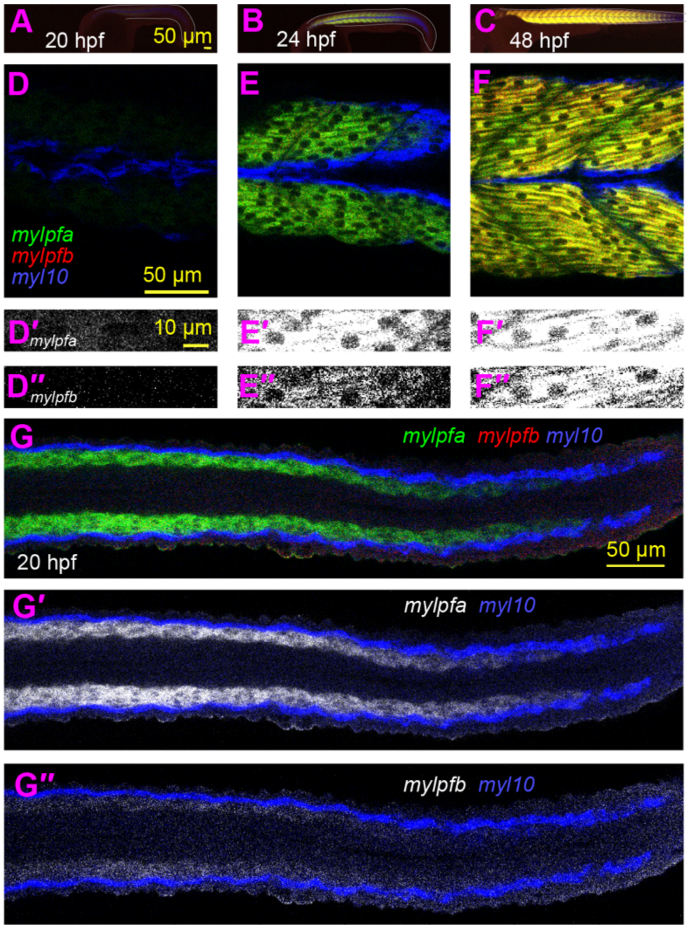

**Figure S3:** Explanation of myofibril bundle width measurements. **(A, B)** Confocal images of 48 hpf embryos labeled for F-actin, MyHC, and Actinin, showing ROI measurements for myofibril bundle width, sampled across the fast-twitch region of somites in a wild-type animal (A) and a *mylpfa*$^{-/-}$ mutant (B). The measures from individual images are averaged and used as one data point for statistical analysis. **(C-D)** Box plots comparing width measurements across genotypes and measurement styles. Average of measurements from the merged image (C) or from the F-actin channel alone (D) appear highly similar. In subsequent analysis, widths were measured using the phalloidin label alone. Comparisons use Student's T-test and matching results are found with Kruskal-Wallis exact test; *** P<0.001.

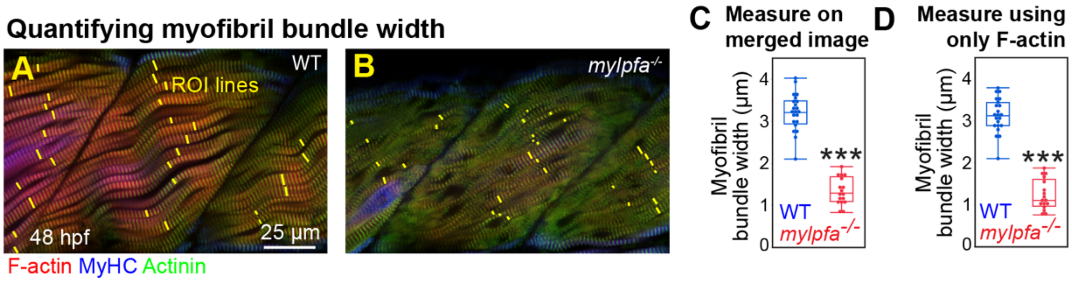

**Figure S4:** Explanation of sarcomeric fraction calculations. **(A, B)** Confocal images of 48 hpf embryos labeled for F-actin, MyHC, and Actinin, showing ROI measurements for sarcomeric fraction, sampled across the shown somites in a wild-type (A) and a *mylpfa*$^{-/-}$ mutant (B). **(C)** Overview of a method to quantify the degree to which markers localize to sarcomeres. 1) ROI lines are drawn throughout the fast-twitch muscle region of a dorsal somite half, with lengths of approximately 15 µm each. The dorsal fast-twitch muscle is used for all measurements of myofibril bundle width in this paper. Examples of wild-type and *mylpfa*$^{-/-}$ measurements are shown at the top of this panel. 2) A FIJI script separates channels and calculates grayscale intensity across the ROI line shown. 3) The periodicity of each ROI is calculated, and repeats are binned by length. 4) Histogram plot of all ROI in one image, here binned to step size 0.05 µm, with peak frequency near the sarcomere length of 1.85 µm. 5) Each 0.05 µm histogram bin is averaged per image, then bootstrap confidence intervals are calculated using variation between images. **(D-G)** Histograms showing the frequency of periodicity in images. Colored lines represent mean values per genotype, semi-transparent colored lines indicate bootstrap confidence intervals, and gray bars indicating the sarcomeric intervals per marker. **(H)** Box plot of sarcomere lengths in the *mylpfa*$^{-/-}$ mutant and their wild-type siblings, showing no change in length, allowing us to use consistent length bins when assessing protein localization to sarcomeres. **(I)** Formula for calculating sarcomeric fraction, which is the ratio of signal in sarcomeric lengths (gray bars) to total localization (Y-axis, 0-4 µm). **(J)** Sarcomeric fractions are shown for F-actin (phalloidin), Z-disk (antibody to Actinin), M-line (antibody to Myomesin), and MyHC (antibody A4.1025) labels. A horizontal brown line shows the fraction predicted by a uniform distribution. For all bar graphs and histogram plots, wild-type data is blue, and the *mylpfa*$^{-/-}$ mutant data is red. Significance thresholds were determined by Kruskal-Wallis exact test (H) or Tukey-Kramer HSD comparisons (J); ** P<0.01, *** P<0.001.

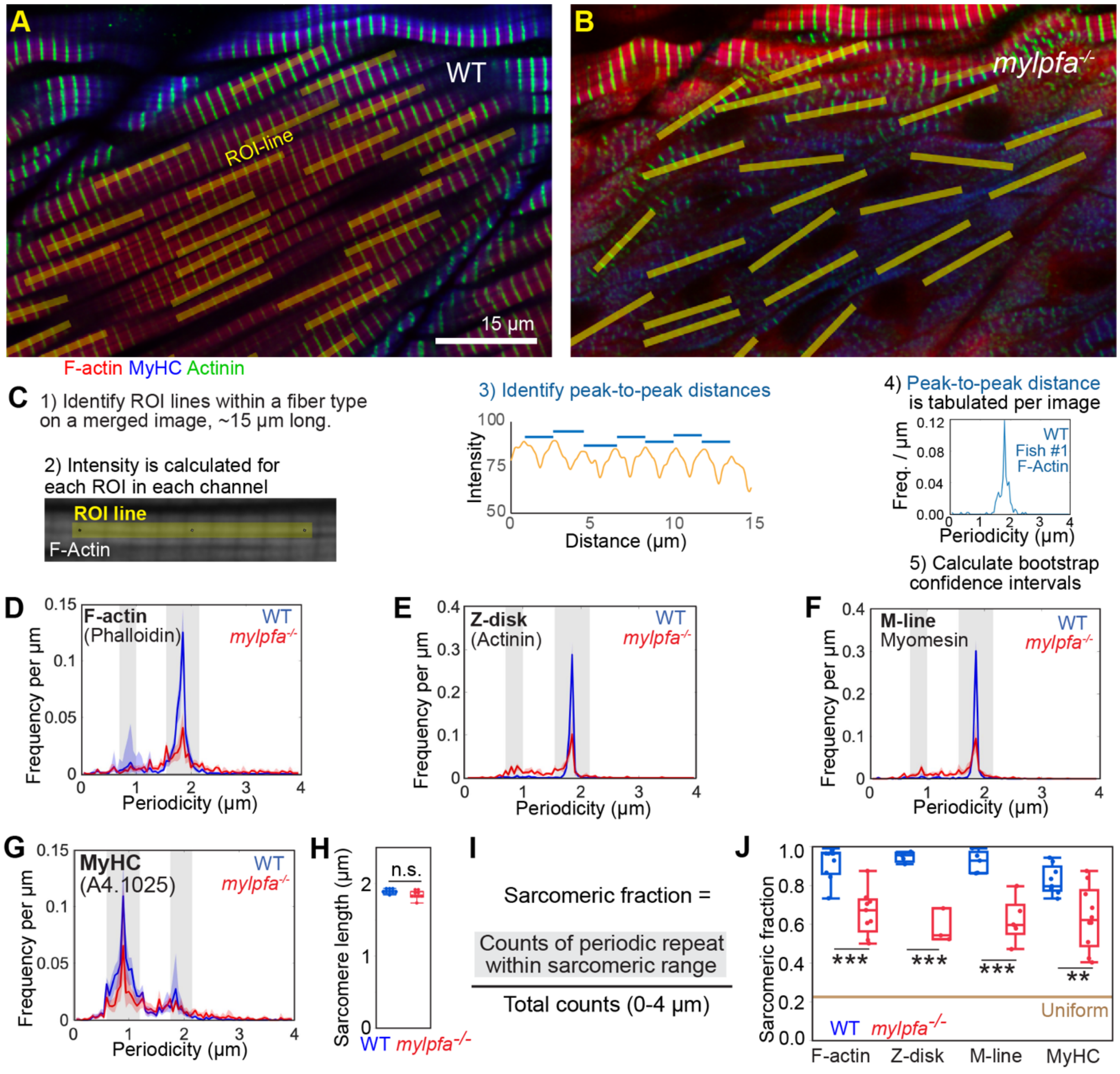

**Figure S5:** <u>The *mylpfa*$^{oz30}$ phenotypes are consistent with those found in the *mylpfa*$^{oz43}$ mutant.</u> **(A)** Illustration of the two *mylpfa* alleles on DNA and **(B)** their effects on the protein's primary sequence. **(C)** Model of Mylpfa protein in complex with a MyHC fragment, generated using Robetta. Color code in the Mylpfa protein shows the portion present in both mutants (green), absent in both mutants (black), and absent only in the *mylpfa*$^{oz43}$ mutant (orange). **(D-F)** Confocal slices through the fast-twitch region of somites labeled with phalloidin at 72 hpf. Compared to wild-type siblings (D), myofibrils are disarrayed in the *mylpfa*$^{oz43}$ mutant (E) and the *mylpfa*$^{oz30}$ mutant (F). Scale bar in D applies to D-F.

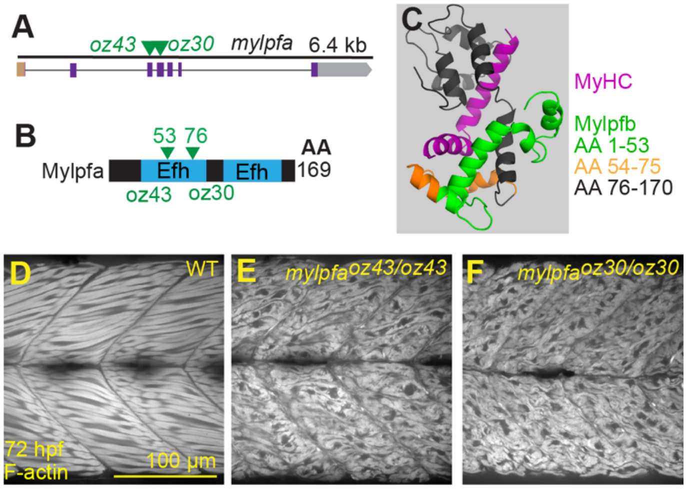

**Figure S6:** <u>Viability and adult phenotypes in the *mylpfa;mylpfb* mutant series.</u> **(A)** Summary of how the number of fish identified during genotyping from adult *mylpfa*$^{+/-}$;*mylpfb*$^{+/-}$ incross compares to expectations. Amongst surviving genotypes there is no significant difference from Mendelian expectation, when the missing genotypes are each considered lethal (Chi$^2$>0.2). By contrast, a nonlethal Mendelian pattern is very unlikely (Chi$^2$<1x10$^{-8}$). All genotypes that are lethal have tattered myofibril bundles and lack high-speed movement at 6 dpf. All surviving genotypes have cohesive myofibril bundles and can move rapidly at 6 dpf. **(B, C)** Images of a wild-type animal (B) and its *mylpfb*$^{-/-}$ sibling (C). **(D, E)** Box plot showing a lack of significant difference in size or weight between these genotypes at 11 months post fertilization. Scalebar in C is for B-C. Significance thresholds are determined by Tukey-Kramer HSD comparisons after one-way ANOVA: n.s. indicates P>0.1. Each point in D and E represents measurement from one animal, which together comprise the statistical N.

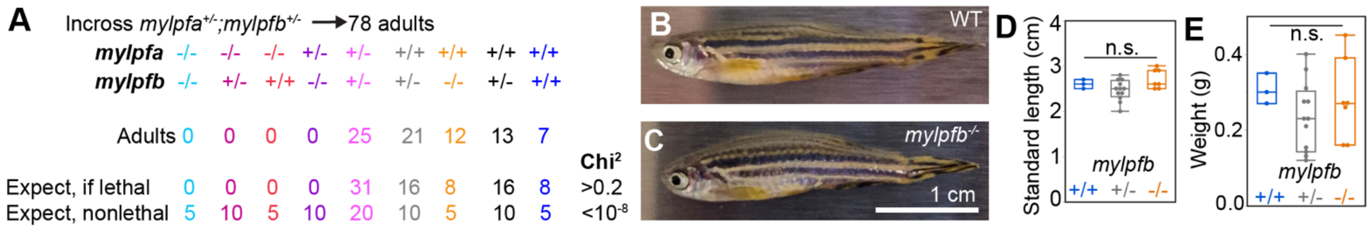

**Figure S7:** <u>Significant increases in slow-twitch speeds are found with strong loss of Mylpf function.</u> **(A-C)** Plot showing the percent of time (Y-axis) spent moving in each speed bin (X-axis) for select genotypes. Speed is binned to every 1 mm/sec (A), every 10 mm/sec (B), or a range of values based on the natural log (C). **(A'-C')** The same data shown normalized to wild-type. We find that the 1 mm/sec speed bins over-segments data at both low and high speed (A, A'). The 10 mm/sec speed bin plot insufficiently samples slow speeds, reducing the 20-40 mm/sec range to just three bins (B, B'). The natural log based bin formula samples movement well across all behavioral ranges (C, C'). The wild-type, *mylpfa*$^{-/-}$, and *mylpfa*$^{-/-}$*;mylpfb*$^{-/-}$ data used for these graphs are also used in Figure 6G-J. Color code, also shown in panel A: WT (blue), *mylpfa*$^{+/-}$*;mylpfb*$^{-/-}$ (purple), *mylpfa*$^{-/-}$ (red), *mylpfa*$^{-/-}$*;mylpfb*$^{+/-}$ (maroon), *mylpfa*$^{-/-}$*;mylpfb*$^{-/-}$ (cyan). The data shown includes all the genotypes from the *mylpfa;mylpfb* double mutant series that lack rapid movements; each shows a significant increase in slow-twitch movement compared to wild-type siblings. Solid lines represent means, shaded lines show bootstrap confidence intervals. Arrows point to an area where all shown mutant genotypes are significantly higher than wild-type siblings in all three binning methods. Y-axis graph ticks shown to indicate linear vs logarithmic scales. X-axis graph ticks indicate each speed bin.

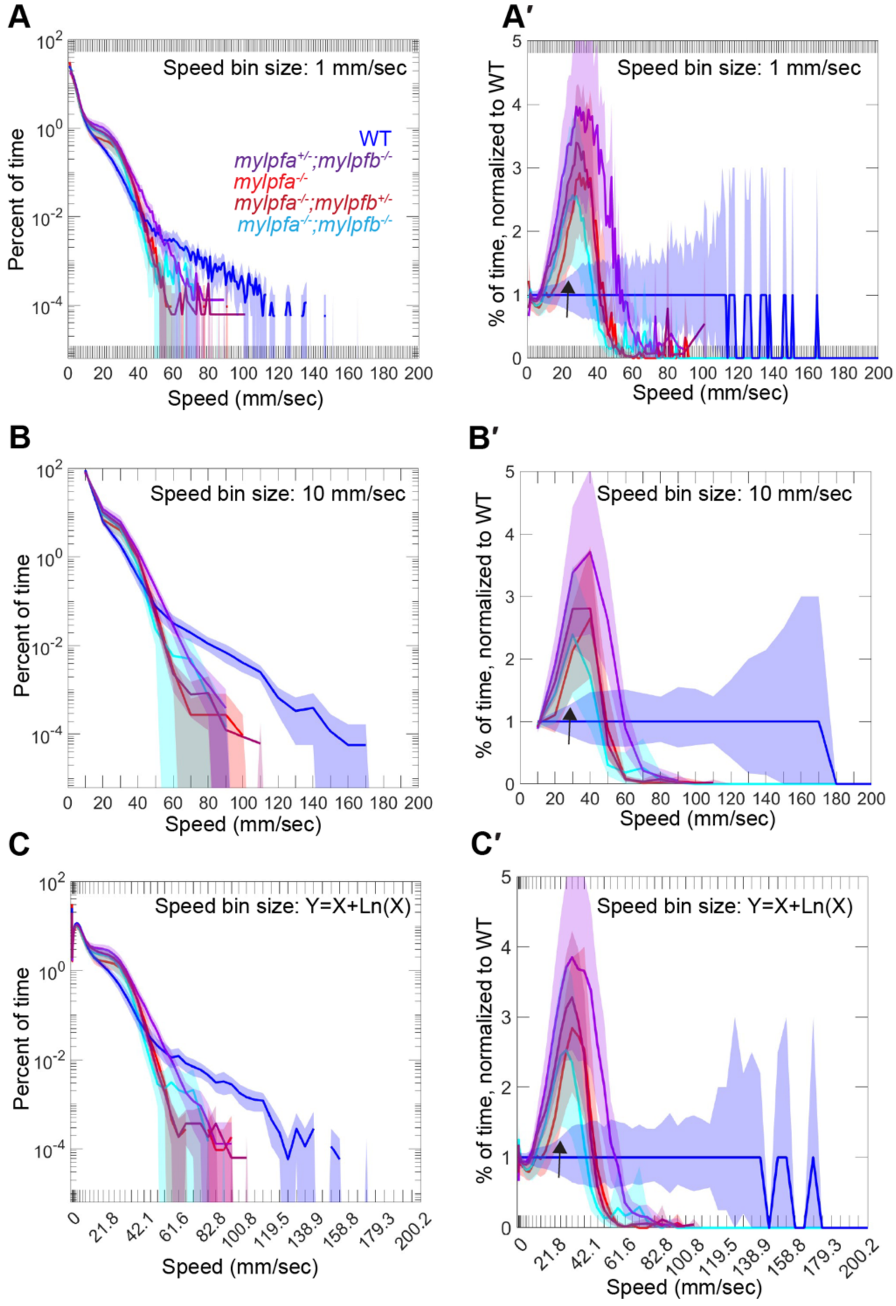

**Figure S8:** Further evidence of slow muscle expansion in the *mylpfa*$^{-/-}$ mutant. **(A, B)** 3D render of confocal stack shows that the wild-type animal (A) has narrower slow muscle fibers than the *mylpfa*$^{-/-}$ mutant (B). The difference is most apparent by the gaps between fibers (pink arrowheads) which almost disappear in the mylpfa mutant. **(C-D″)** Cross view of somite labeled for phalloidin and F59, showing that the F59 label becomes brighter and wider in the *mylpfa*$^{-/-}$ mutant.

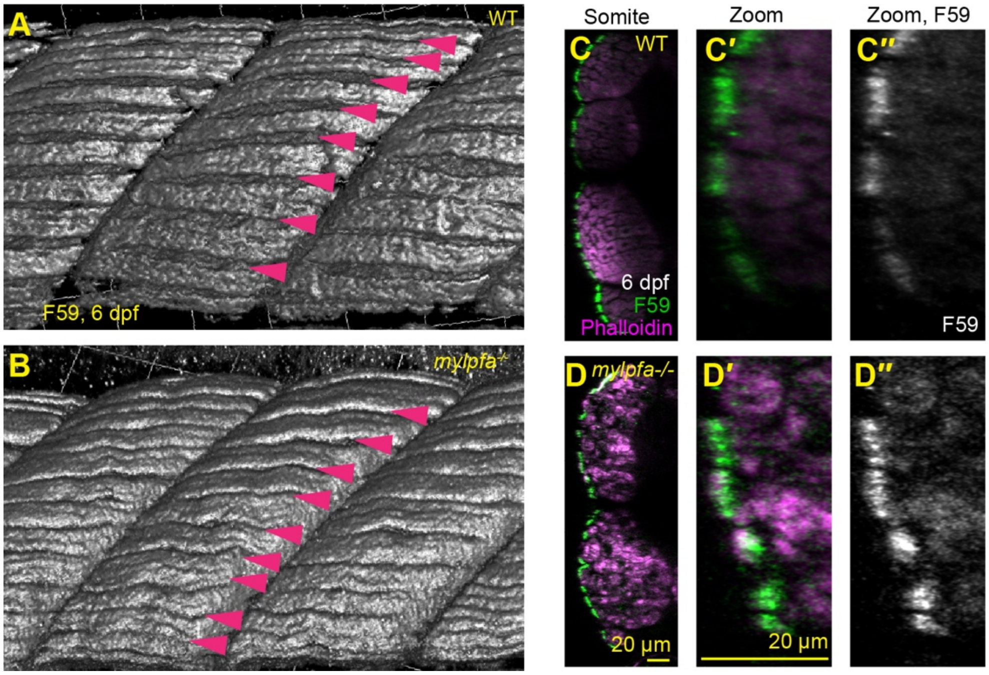

**Figure S9:** Mosaic expression of either *mylpfb-GFP* or *mylpfa-GFP* can rescue the *mylpfa*$^{-/-}$ myofibrils. **(A-D)** Images of phalloidin labeled animals at 72 hpf showing mosaic expression of the *mylpfa:mylpfa-GFP* (A, B) or the *mylpfa:mylpfb-GFP* transgene (C, D). **(E)** Box plots showing the fraction of F-actin localized to sarcomeres, calculated in non-transgenic muscle fibers and GFP+ muscle fibers within the same mosaic animals. **(F)** Box plots showing the sarcomeric fraction for GFP. **(G)** Scatterplot showing the correlation between GFP brightness and myofibril width in *mylpfa*$^{-/-}$ mutant animals carrying *mylpfa:mylpfa-GFP* or *mylpfa:mylpfb-GFP* transgenes. Linear correlates for the two transgenes have overlapping confidence intervals. In mosaic measurements (E, F), each GFP+ point represents the mean of measurements from fibers in an individual fish with GFP brightness above 10. The GFP- points were measured in GFP-negative fibers of the same animals. The transgenic measurements (G) compare measurements of transgenic animals versus non-transgenic siblings. The vertical brown line indicates animals classified as transgenic vs. non-transgenic. Significance: n.s. is P>0.1, *** indicates P<0.001 as determined by Tukey-Kramer HSD comparisons after one-way ANOVA.

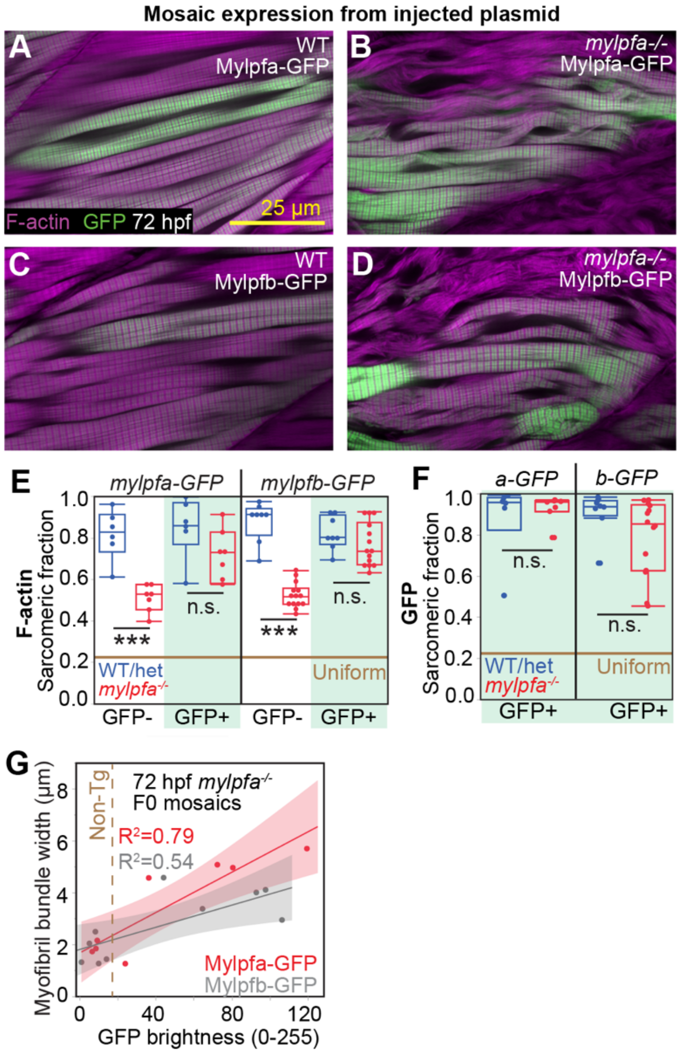

**Figure S10:** <u>The *mylpfa:mylpfa-GFP* transgene restores muscle morphology in both fast-twitch and slow-twitch muscle at 6 dpf.</u> **(A)** Measurements of myofibril bundle width taken in sagittal-view of fast-twitch muscle at 6 dpf. **(B-E)** Examples of cross-sectional images, reconstructed from confocal stacks with insets from boxed areas shown beneath the overview. The myofibril-free central region of the muscle fiber's central cytoplasm (Arrowhead) typically expands in the non-transgenic *mylpfa*$^{-/-}$ mutant (C) but is reduced in the *mylpfa*$^{-/-}$ mutant carrying the transgene (E). **(F)** Measurement of myofibril cross sectional area within the slow-twitch region shows an expansion in the *mylpfa*$^{-/-}$ single mutant and rescue to the smaller wild-type size in the *mylpfa*$^{-/-}$ mutants that express the *mylpfa:mylpfa-GFP* transgene. Significance in F is determined using Tukey-Kramer comparisons after one-way ANOVA, significance shown as n.s. of P>0.1 and ** P<0.01. Points in panels A and F represent statistical N, which is the mean of measurements in individual animals. Both 10 µm scalebars in B are for B-E.

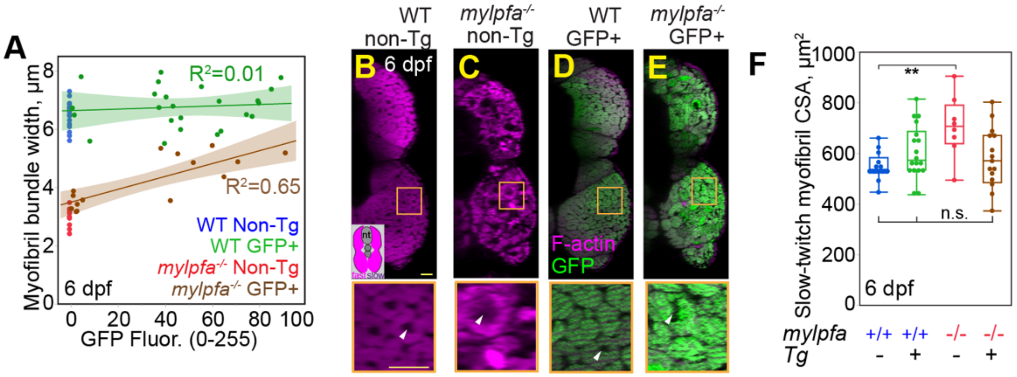

**Figure S11**: Further quantification showing that the G163S variant is unable to promote myofibril formation. **(A)** Statistical comparisons of myofibril bundle width in animals mosaically expressing wild-type or G163S MYLPF variant protein; the non-Tg group represents the non-transgenic fibers from these animals. **(B)** Box plot of the sarcomeric fraction for F-actin in the same animals. **(C, D)** Histograms showing the frequency of periodicity in the 2 dpf images, calculated as explained in Figure S4. Colored lines represent mean values per genotype, semi-transparent colored lines indicate bootstrap confidence intervals, and gray bars indicating the sarcomeric intervals per marker. Myofibers that express the MYLPF-GFP protein in the *mylpfa*$^{-/-}$ mutant show a significant increase in the sarcomeric signal compared to GFP-negative (GFP<10) myofibers in the same animals. By contrast, in *mylpfa*$^{-/-}$ mutant animals expressing the G163S protein variant, GFP+ myofibers do not significantly differ from GFP-negative fast-twitch muscle fibers. Panels A-D represent all animals shown in Figure 9, plus two more repeats that had uneven GFP settings which were excluded from brightness analysis (Figure 9). Each GFP+ point (WT and G163S) represents the mean of measurements from fibers in an individual fish with GFP brightness >10. The GFP- points were measured in GFP- fibers of the same animals. Significance in A and B is determined by Tukey-Kramer HSD comparisons after ANOVA: n.s. is P>0.1, * indicates P<0.05, and *** indicates P<0.001.

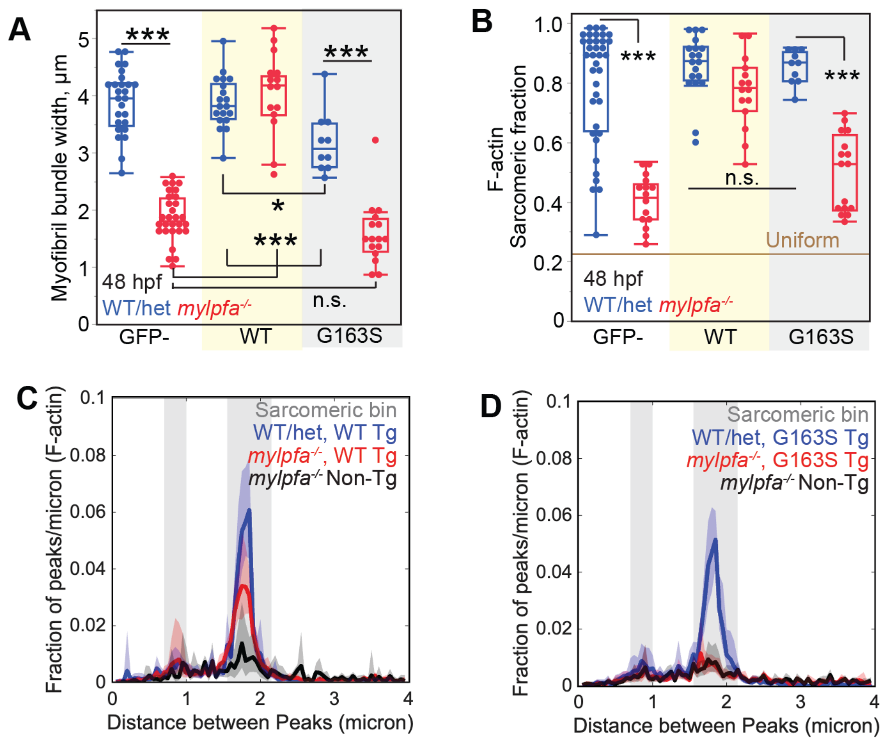

**Figure S12:** Explanation of data mergers for final model. We wished to compare the effect of Mylpf dosage on muscle phenotype across experiments, timepoints, and analytic methods. **(A-A")** Graphs of transgenic analysis that were conducted at 2 and 6 dpf, leading to different width ranges by stage. The shown panels display data from Figures 7I (A), S10A (A'), and 8B (A"). **(B)** We developed a simple set of formulas to scale these datasets along shared axes, using the 2:1 ratio between Mylpfa and Mylpfb at 48 hpf. The myofibril bundle widths are converted to an approximately 0-1 scale by dividing by the WT mean per experiment. No transformation is needed for the sarcomeric fraction, which is already calculated on a 0-1 scale. **(C)** After data is processed through these formulas, they can be overlaid; this merger is reproduced in Figure 8F. We find no significant differences in trendlines between the Mylpfa-GFP, Mylpfb-GFP, or MYLPF-GFP transgenic analysis in *mylpfa*$^{-/-}$, nor in WT. **(D)** We also merged the *mylpfa;mylpfb* mutant analysis at 1, 2, 3, and 6 dpf, using data shown in Figure 5A, B, C, and D respectively. **(D')** A subset of this data showing the two genotypes with lowest Mylpf dose at 2 and 6 dpf. This data is merged with transgenic data to form a lower asymptote during Hill regression. **(E)** Plot of the low-dose genotypes merged with transgenic data (GFP>10) and resulting regression model. Both the Hill slope and the EC50 significantly differ from the regression model built using the *mylpfa;mylpfb* mutant series. **(F)** A similar model can be produced using the F-actin sarcomeric fraction. With this measure, we find no significant difference in the Hill slope between the transgenic data and the mutant data. The color code in C-F matches that shown in A-A''' and D. Panels D and E reproduce figure 9A and B respectively, shown here to give context about their construction. Asterisks indicate difference between the transgenic Hill regressions and the Hill regressions from *mylpfa;mylpfb* mutant series; * indicates P<0.05 and *** P<0.001.

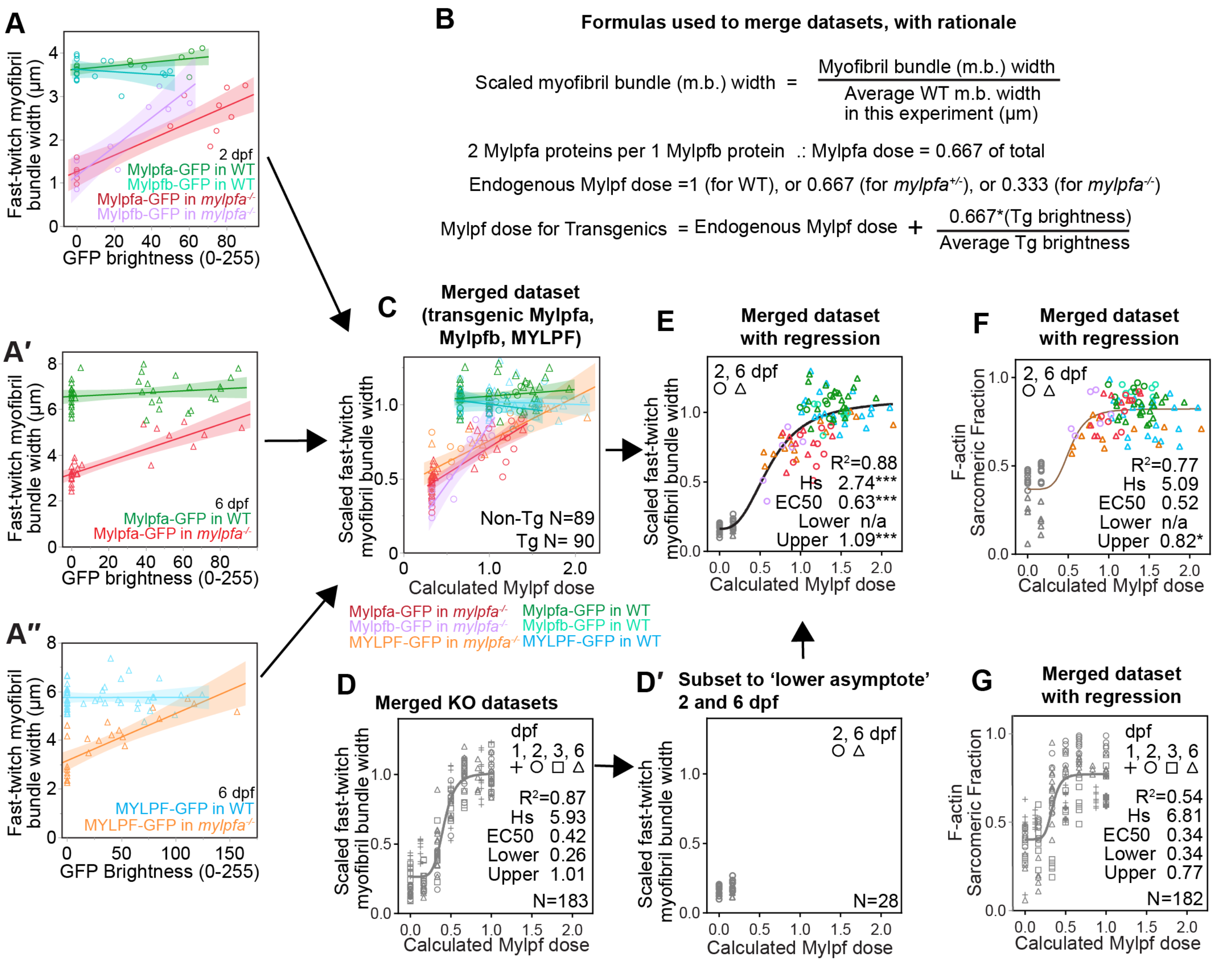

**Video 1:** Confocal stacks showing differences in muscle structure between wild-type and *mylpfa*$^{-/-}$ sibling animals. Animals were fixed at 48 hpf then labeled for F-actin (red), Actinin (green), and MyHC (blue). The video contains lateral slow-twitch fibers and the superficial layers of fast-twitch fibers layers of a dorsal half-somite. Fast-twitch muscle fibers in the centered somite are indicated by white ovals. Scalebar in the wild-type stack also applies to the *mylpfa*$^{-/-}$ mutant stack.

**Video 2:** The *mylpfa*$^{-/-}$ mutant shows an impaired escape response. Imaging at 240 frames/second, showing 100 msec after contact with fishing line in the wild-type and the *mylpfa*$^{-/-}$ mutant. Both animals respond to the prodding; however, the *mylpfa*$^{-/-}$ mutant escapes at slower speed due to smaller tail undulations. Figure 6A, B shows a projection of frames from this video.

**Tables:**

**Table S1:** Calculating the predicted doses for Mylpf in the *mylpfa;mylpfb* mutant series. "Genotype shorthand" offers an abbreviated way to refer to each genotype. Color code matches the genotypic color code used throughout the manuscript. For each genotype, we calculated predicted Mylpf abundance using a 1:1 Mylpfa to Mylpfb ratio (gene dose), a 7:1 ratio (24 hpf protein dose), and a 2:1 ratio (48hpf+ protein dose). The western blot (Observed Mylpf) shows a near-linear trend between 100% (wild-type) and 0% (double mutant). Observed Mylpf abundance does not substantially differ from 48 hpf expectations in any genotype. By contrast, the observed MyHC abundance is even across genotypes.

| Genotype Shorthand | Genotype | | 1:1 Formula | Calculated Mylpf (1:1) | 7:1 Formula | Calculated Mylpf (7:1) | 2:1 Formula | Calculated Mylpf (2:1) | Observed Mylpf | Observed MyHC |
|---|---|---|---|---|---|---|---|---|---|---|
| | *mylpfa* | *mylpfb* | | | | | | | | |
| **WT** | +/+ | +/+ | $\frac{1+1+1+1}{4}$ | 1.00 | $\frac{7+7+1+1}{16}$ | 1.000 | $\frac{2+2+1+1}{6}$ | 1.000 | 1.000 | 1.000 |
| **Bhet** | +/+ | +/- | $\frac{1+1+1+0}{4}$ | 0.75 | $\frac{7+7+1+0}{16}$ | 0.938 | $\frac{2+2+1+0}{6}$ | 0.833 | 0.883 | 0.926 |
| **Bmut** | +/+ | -/- | $\frac{1+1+0+0}{4}$ | 0.50 | $\frac{7+7+0+0}{16}$ | 0.875 | $\frac{2+2+0+0}{6}$ | 0.667 | 0.643 | 1.031 |
| **Ahet** | +/- | +/+ | $\frac{1+0+1+1}{4}$ | 0.75 | $\frac{7+0+1+1}{16}$ | 0.563 | $\frac{2+0+1+1}{6}$ | 0.667 | 0.825 | 1.322 |
| **AhetBhet** | +/- | +/- | $\frac{1+0+1+0}{4}$ | 0.50 | $\frac{7+0+1+0}{16}$ | 0.500 | $\frac{2+0+1+0}{6}$ | 0.500 | 0.615 | 1.198 |
| **AhetBmut** | +/- | -/- | $\frac{1+0+0+0}{4}$ | 0.25 | $\frac{7+0+0+0}{16}$ | 0.438 | $\frac{2+0+0+0}{6}$ | 0.333 | 0.398 | 0.916 |
| **Amut** | -/- | +/+ | $\frac{0+0+1+1}{4}$ | 0.50 | $\frac{0+0+1+1}{16}$ | 0.125 | $\frac{0+0+1+1}{6}$ | 0.333 | 0.278 | 0.844 |
| **AmutBhet** | -/- | +/- | $\frac{0+0+1+0}{4}$ | 0.25 | $\frac{0+0+1+0}{16}$ | 0.063 | $\frac{0+0+1+0}{6}$ | 0.167 | 0.105 | 0.823 |
| **Double** | -/- | -/- | $\frac{0+0+0+0}{4}$ | 0.00 | $\frac{0+0+0+0}{16}$ | 0.000 | $\frac{0+0+0+0}{6}$ | 0.000 | 0.000 | 0.910 |

**Table S2:** Summary of statistical tests for regression using the Hill equation. The phenotypic data from the *mylpfa;mylpfb* mutant series, shown in Figure 5, was run through regression models using the Hill equation with different levels of *mylpfa* and *mylpfb* skewing. Calculations are shown for the myofibril bundle width (width), F-actin sarcomeric fraction (F-actin s.f.), and myosin heavy chain sarcomeric fraction (MyHC s.f.) In all cases, direct allele count (the gene dose: 1 *mylpfa* allele per 1 *mylpfb* allele) provides low predictive value ($R^2$) compared to mRNA and protein ratios. These mRNA and Protein doses are calculated using the 7:1 Mylpfa to Mylpfb ratio observed at 1 dpf and a 2:1 Mylpfa to Mylpfb ratio for 2 dpf onwards. Hill slopes that are calculated by this mildly skewed ratio are more consistent than those calculated using the highly skewed 7 to 1 ratio seen at 1 dpf. All valid models give Hill slopes larger than one. A gray box indicates an invalid regression model. Bolding indicates column headers. Thick borders indicate data shown in Figure 5.

| | | **26 hpf** | **48 hpf** | | | | **72 hpf** | **6 dpf** |
|---|---|---|---|---|---|---|---|---|
| | | **Width** | **Width** | **F-actin s.f.** | **MyHC s.f.** | **Colocalization** | **Width** | **Width** |
| **1:1** | **Hs** | | 5.28 | 5.52 | 16.48 | 3.56 | 4.47 | 1.76 |
| | **EC50** | N/A | 0.43 | 0.39 | 0.46 | 0.40 | 0.46 | 0.60 |
| | **$R^2$** | | 0.71 | 0.80 | 0.74 | 0.52 | 0.69 | 0.57 |
| **2:1** | **Hs** | 7.26 | 6.21 | 9.39 | 11.35 | 6.84 | 4.46 | 4.18 |
| | **EC50** | 0.41 | 0.41 | 0.34 | 0.35 | 0.43 | 0.43 | 0.40 |
| | **$R^2$** | 0.84 | 0.95 | 0.94 | 0.92 | 0.78 | 0.88 | 0.89 |
| **7:1** | **Hs** | 1.93 | 1.57 | 3.18 | 16.29 | 1.28 | 1.28 | 1.63 |
| | **EC50** | 0.30 | 0.30 | 0.13 | 0.13 | 1.28 | 0.39 | 0.41 |
| | **$R^2$** | 0.84 | 0.95 | 0.94 | 0.92 | 0.79 | 0.88 | 0.90 |
